# Close relatives of MERS-CoV in bats use ACE2 as their functional receptors

**DOI:** 10.1101/2022.01.24.477490

**Authors:** Qing Xiong, Lei Cao, Chengbao Ma, Chen Liu, Junyu Si, Peng Liu, Mengxue Gu, Chunli Wang, Lulu Shi, Fei Tong, Meiling Huang, Jing Li, Chufeng Zhao, Chao Shen, Yu Chen, Huabin Zhao, Ke Lan, Xiangxi Wang, Huan Yan

**Affiliations:** State Key Laboratory of Virology, Institute for Vaccine Research and Modern Virology Research Center, College of Life Sciences, Wuhan University, Wuhan, Hubei, 430072, China; CAS Key Laboratory of Infection and Immunity, National Laboratory of Macromolecules, Institute of Biophysics, Chinese Academy of Sciences, Beijing, 100101, China; University of Chinese Academy of Sciences, Beijing, 100049, China; Department of Ecology, Tibetan Centre for Ecology and Conservation at WHU-TU, Hubei Key Laboratory of Cell Homeostasis, College of Life Sciences, Wuhan University, Wuhan 430072, China

**Keywords:** NeoCoV, PDF-2180-CoV, MERS-CoV, bat merbecovirus, ACE2, DPP4

## Abstract

Middle East Respiratory Syndrome coronavirus (MERS-CoV) and several bat coronaviruses employ Dipeptidyl peptidase-4 (DPP4) as their functional receptors^1–4^. However, the receptor for NeoCoV, the closest MERS-CoV relative yet discovered in bats, remains enigmatic^5^. In this study, we unexpectedly found that NeoCoV and its close relative, PDF-2180-CoV, can efficiently use some types of bat Angiotensin-converting enzyme 2 (ACE2) and, less favorably, human ACE2 for entry. The two viruses use their spikes’ S1 subunit carboxyl-terminal domains (S1-CTD) for high-affinity and species-specific ACE2 binding. Cryo-electron microscopy analysis revealed a novel coronavirus-ACE2 binding interface and a protein-glycan interaction, distinct from other known ACE2-using viruses. We identified a molecular determinant close to the viral binding interface that restricts human ACE2 from supporting NeoCoV infection, especially around residue Asp338. Conversely, NeoCoV efficiently infects human ACE2 expressing cells after a T510F mutation on the receptor-binding motif (RBM). Notably, the infection could not be cross-neutralized by antibodies targeting SARS-CoV-2 or MERS-CoV. Our study demonstrates the first case of ACE2 usage in MERS-related viruses, shedding light on a potential bio-safety threat of the human emergence of an ACE2 using “MERS-CoV-2” with both high fatality and transmission rate.

## Introduction

Coronaviruses (CoVs) are a large family of enveloped positive-strand RNA viruses classified into four genera: Alpha-, Beta-, Gamma- and Delta-CoV. Generally, Alpha and Beta-CoV can infect mammals such as bats and humans, while Gamma- and Delta-CoV mainly infect birds, occasionally mammals^6–8^. It is thought that the origins of most coronaviruses infecting humans can be traced back to their close relatives in bats, the most important animal reservoir of mammalian coronaviruses ^9, 10^. Coronaviruses are well recognized for their recombination and host-jumping ability, which has led to the three major outbreaks in the past two decades caused by SARS-CoV, MERS-CoV, and the most recent SARS-CoV-2, respectively^11–14^.

MERS-CoV belongs to the linage C of Beta-CoV (Merbecoviruses), which poses a great threat considering its high case-fatality rate of approximately 35%^15^. Merbecoviruses have also been found in several animal species, including camels, hedgehogs, and bats. Although camels are confirmed intermediate hosts of the MERS-CoV, bats, especially species in the family of *Vespertilionidae*, are widely considered to be the evolutionary source of MERS-CoV or its immediate ancestor^16^.

Specific receptor recognition of coronaviruses is usually determined by the receptor-binding domains (RBDs) on the carboxyl-terminus of the S1 subunit (S1-CTD) of the spike proteins^17^. Among the four well-characterized coronavirus receptors, three are S1-CTD binding ectopeptidases, including ACE2, DPP4, and aminopeptidase N (APN)^1, 18, 19^. By contrast, the fourth receptor, antigen-related cell adhesion molecule 1(CEACAM1a), interacts with the amino-terminal domain (NTD) of the spike S1 subunit of the murine hepatitis virus^20, 21^. Interestingly, the same receptor can be shared by distantly related coronaviruses with structurally distinct RBDs. For example, the NL63-CoV (an alpha-CoV) uses ACE2 as an entry receptor widely used by many sarbecoviruses (beta-CoV linage B)^22^. A similar phenotype of cross-genera receptor usage has also been found in APN, which is shared by many alpha-CoVs and a delta-CoV (PDCoV)^7^. In comparison, DPP4 usage has only been found in merbecoviruses (beta-CoV linage C) such as HKU4, HKU25, and related strains^2–4^.

Intriguingly, many other merbecoviruses do not use DPP4 for entry and their receptor usage remains elusive, such as bat coronaviruses NeoCoV, PDF-2180-CoV, HKU5-CoV, and hedgehog coronaviruses EriCoV-HKU31^5, 23–25^. Among them, the NeoCoV, infecting *Neoromicia capensis* in South Africa, represents a bat merbecovirus that happens to be the closest relative of MERS-CoV (85% identity at the whole genome level)^26, 27^. PDF-2180-CoV, another coronavirus most closely related to NeoCoV, infects *Pipisrellus hesperidus* native to Southwest Uganda^23, 28^. Indeed, NeoCoV and PDF-2180-CoV share sufficient similarity with MERS-CoV across most of the genome, rendering them taxonomically the same viral species^27, 29^. However, their S1 subunits are highly divergent compared with MERS-CoV (around 43-45% amino acid similarity), in agreement with their different receptor preference^23^.

In this study, we unexpectedly found that NeoCoV and PDF-2180-CoV use bat ACE2 as their functional receptor. The cryo-EM structure of NeoCoV RBD bound with the ACE2 protein from *Pipistrellus pipistrellus* revealed a novel ACE2 interaction mode that is distinct from how human ACE2 (hACE2) interacts with the RBDs from SARS-CoV-2 or NL63. Although NeoCoV and PDF-2180-CoV cannot efficiently use hACE2 based on their current sequences, the spillover events of this group of viruses should be closely monitored, considering their human emergence potential after gaining fitness through antigenic drift.

## Results

### Evidence of ACE2 usage

To shed light on the relationship between merbecoviruses, especially NeoCoV and PDF-2180-CoV, we conducted a phylogenetic analysis of the sequences of a list of human and animal coronaviruses. Maximum likelihood phylogenetic reconstructions based on complete genome sequences showed that NeoCoV and PDF-2180-CoV formed sister clade with MERS-CoV (Fig. 1a). In comparison, the phylogenetic tree based on amino acid sequences of the S1 subunit demonstrated that NeoCoV and PDF-2180-CoV showed a divergent relationship with MERS-CoV but are closely related to the hedgehog coronaviruses (EriCoVs) (Fig. 1b). A sequence similarity plot analysis (Simplot) queried by MERS-CoV highlighted a more divergent region encoding S1 for NeoCoV and PDF-2180-CoV compared with HKU4-CoV (Fig. 1c). We first tested whether human DPP4 (hDPP4) could support the infection of several merbecoviruses through a pseudovirus entry assay^30^. The result revealed that only MERS-CoV and HKU4-CoV showed significantly enhanced infection of 293T-hDPP4. Unexpectedly, we detected a significant increase of entry of NeoCoV and PDF-2180-CoV in 293T-hACE2 but not 293T-hAPN, both of which are initially set up as negative controls (Fig. 1d, Extended Data Fig.1).

**Fig. 1.**
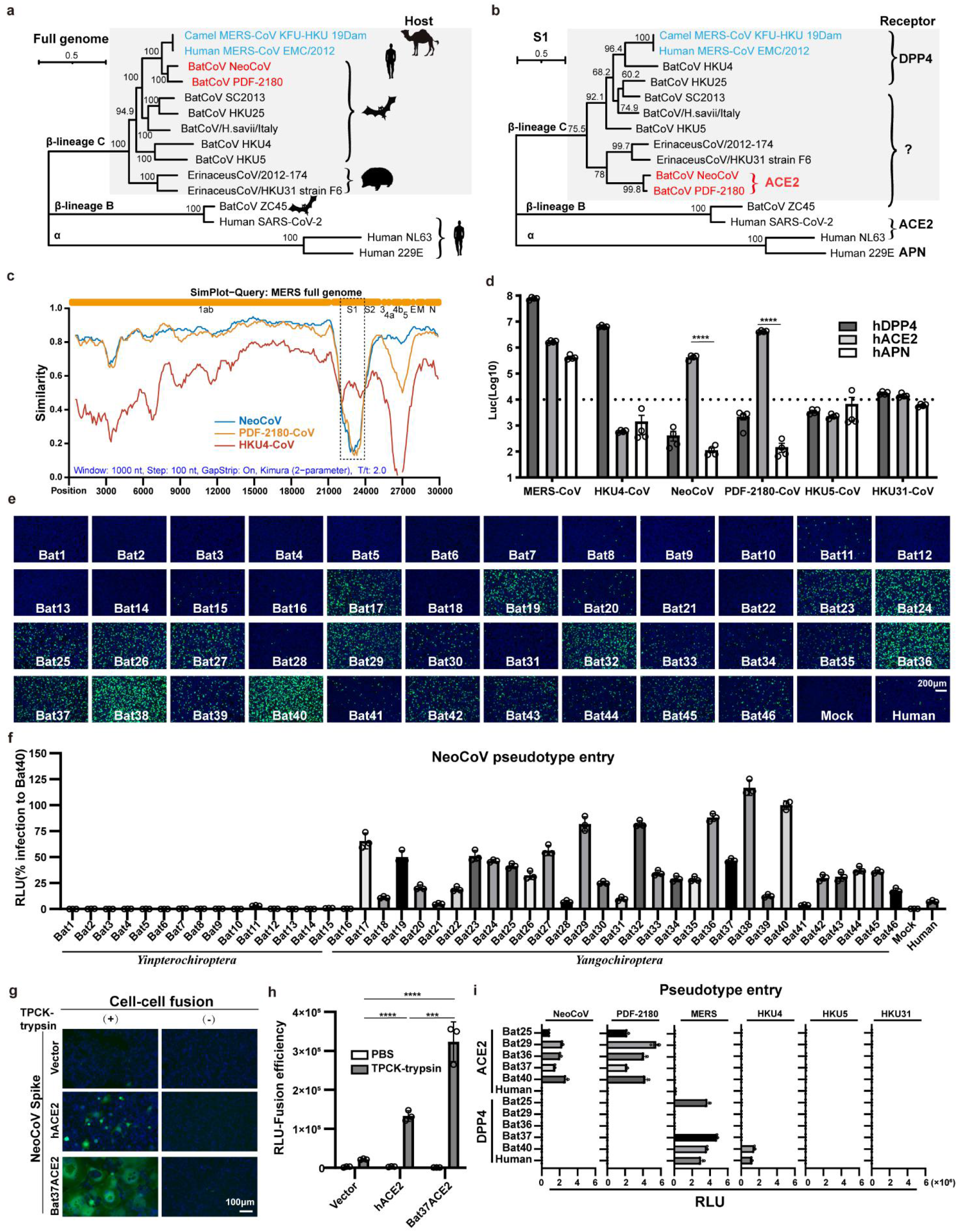
A clade of bat merbecoviruses can use ACE2 but not DPP4 for efficient entry. **a-b,** Phylogenetical analysis of merbecoviruses (gray) based on whole genomic sequences (**a**) and S1 amino acid sequences (**b**). NL63 and 229E were set as outgroups. Hosts and receptor usage were indicated. **c,** Simplot analysis showing the whole genome similarity of three merbecoviruses compared with MERS-CoV. The regions that encode MERS-CoV proteins were indicated on the top. Dashed box: S1 divergent region. **d**, Entry efficiency of six merbecoviruses in 293T cells stably expressing hACE2, hDPP4, or hAPN. **e-f,** Entry efficiency of NeoCoV in cells expressing ACE2 from different bats. EGFP intensity (**e**); firefly luciferase activity (**f**). **g-h,** Cell-cell fusion assay based on dual-split proteins showing the NeoCoV spike protein mediated fusion in BHK-21 cells expressing indicated receptors. EGFP intensity (**g**), live-cell Renilla luciferase activity (**h**). **i,** Entry efficiency of six merbecoviruses in 293T cells stably expressing the indicated bat ACE2 or DPP4. Mean±SEM for **d**, **i;** Mean±SD for **f**, and **h**.(n=3). RLU: relative light unit.

To further validate the possibility of more efficient usage of bat ACE2, we screened a bat ACE2 cell library individually expressing ACE2 orthologs from 46 species across the bat phylogeny, as described in our previous study^31^(Extended Data Figs.2-3, Supplementary Table 1). Interestingly, NeoCoV and PDF-2180-CoV, but not HKU4-CoV or HKU5-CoV, showed efficient entry in cells expressing ACE2 from most bat species belonging to *Vespertilionidae* (vesper bats). In contrast, no entry or very limited entry in cells expressing ACE2 of humans or bats from the *Yinpterochiroptera* group (Fig. 1e-f, Extended Data Fig.4). Consistent with the previous reports, the infection of NeoCoV and PDF-2180-CoV could be remarkably enhanced by an exogenous trypsin treatment^28^(Extended Data Fig.5). As indicated by the dual split protein (DSP)-based fusion assay ^32^, Bat37ACE2 triggers more efficient cell-cell membrane fusion than hACE2 in the presence of NeoCoV spike protein expression (Fig. 1g-h). Notably, the failure of the human or hedgehog ACE2 to support entry of EriCoV-HKU31 indicates that these viruses have a different receptor usage (Extended Data Fig.6). In agreement with a previous study^23, 28^, our results against the possibility that bat DPP4 act as a receptor for NeoCoV and PDF-2180-CoV, as none of the tested DPP4 orthologs, from the vesper bats whose ACE2 are highly efficient in supporting vial entry, could support a detectable entry of NeoCoV and PDF-2180-CoV (Fig. 1i, Extended Data Fig.7). Infection assays were also conducted using several other cell types from different species, including a bat cell line Tb 1 Lu, ectopically expressing ACE2 or DPP4 from Bat40 (*Antrozous pallidus*), and each test yielded similar results (Extended Data Fig.8).

### S1-CTD mediated species-specific binding

The inability of NeoCoV and PDF-2180-CoV to use DPP4 is consistent with their highly divergent S1-CTD sequence compared with the MERS-CoV and HKU4-CoV. We produced S1-CTD-hFc proteins (putative RBD fused to human IgG Fc domain) to verify whether their S1-CTDs are responsible for ACE2 receptor binding. The live-cell binding assay based on cells expressing various bat ACE2 showed a species-specific utilization pattern in agreement with the results of the pseudovirus entry assays (Fig. 2a). The specific binding of several representative bat ACE2 was also verified by flow-cytometry (Fig. 2b). We further determined the binding affinity by Bio-Layer Interferometry (BLI) analysis. The results indicated that both viruses bind to the ACE2 from *Pipistrellus pipistrellus* (Bat37) with the highest affinity (KD=1.98nM for NeoCoV and 1.29 nM for PDF-2180-CoV). In contrast, their affinities for hACE2 were below the detection limit of our BLI analysis (Fig. 2c, Extended Data Fig.9). An enzyme-linked immunosorbent assay (ELISA) also demonstrated the strong binding between NeoCoV/PDF-2180-CoV S1-CTDs and Bat37ACE2, but not hACE2 (Fig. 2d). Notably, as the ACE2 sequences of the hosts of NeoCoV and PDF-2180-CoV are unknown, Bat37 represents the closest relative of the host of PDF-2180-CoV (*Pipisrellus hesperidus*) in our study. The binding affinity was further verified by competitive neutralization assays using soluble ACE2-ectodomain proteins or viral S1-CTD-hFc proteins. Again, the soluble Bat37ACE2 showed the highest activity to neutralize viral infection caused by both viruses (Fig. 2e-f). Moreover, NeoCoV-S1-CTD-hFc could also potently neutralize NeoCoV and PDF-2180-CoV infections of cells expressing Bat37ACE2 (Fig. 2g). We further demonstrated the pivotal role of S1-CTD in receptor usage by constructing chimeric viruses and testing them for altered receptor usage. As expected, batACE2 usage was changed to hDPP4 usage for a chimeric NeoCoV with CTD, but not NTD, sequences replaced by its MERS-CoV counterpart (Fig. 2h). These results confirmed that S1-CTD of NeoCoV and PDF-2180-CoV are RBDs for their species-specific interaction with ACE2.

**Fig. 2.**
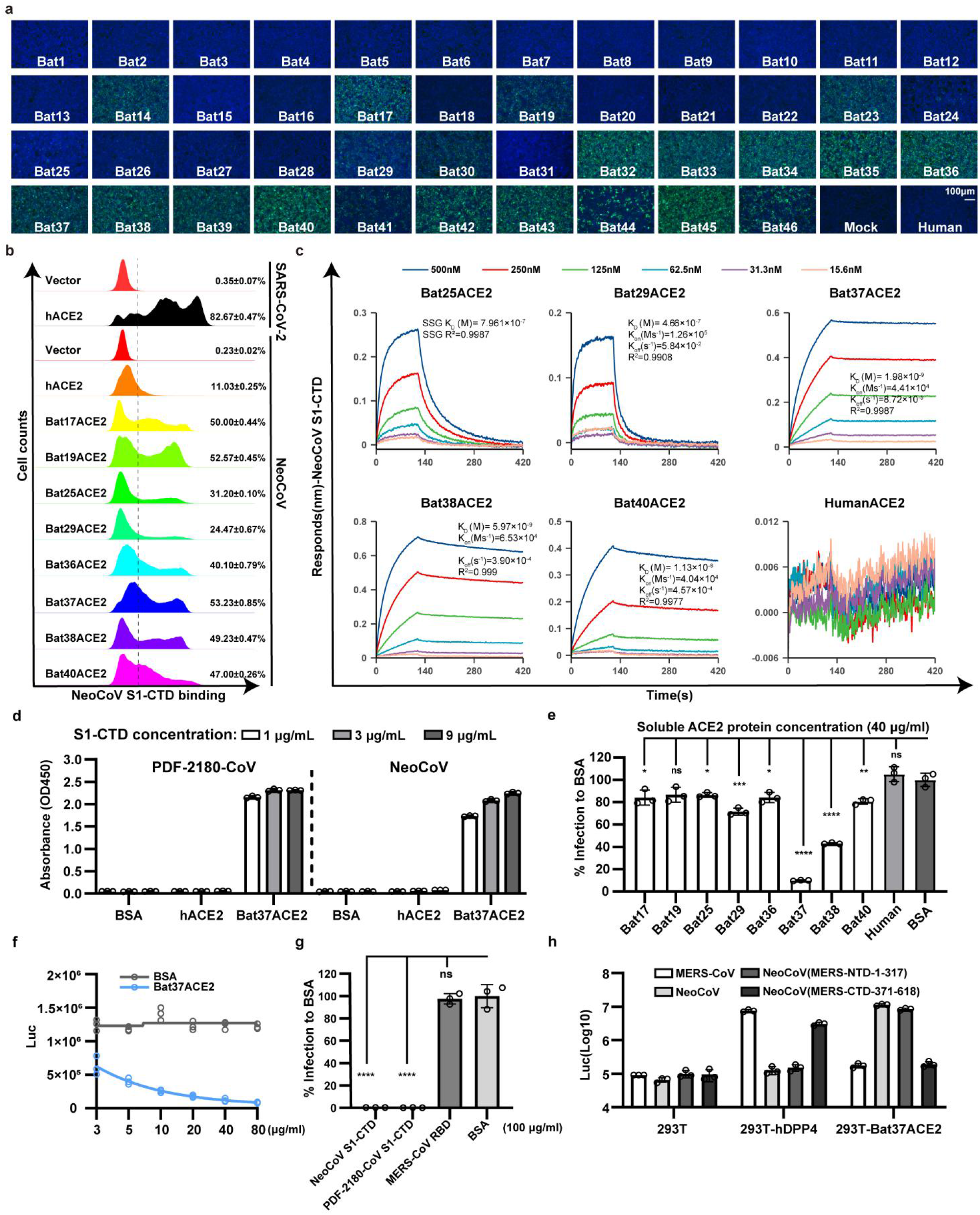
S1-CTD of NeoCoV and PDF-2180-CoV was required for species-specific ACE2 binding. **a,** Binding of NeoCoV-S1-CTD-hFc with 293T bat ACE2 cells via immunofluorescence detecting the hFc. **b,** Flow cytometry analysis of NeoCoV-S1-CTD-hFc binding with 293T cells expressing the indicated ACE2. The positive ratio was indicated based on the threshold (dash line). **c,** BLI assays analyzing the binding kinetics between NeoCoV-S1-CTD-hFc with selected ACE2-ecto proteins. **d,** ELISA assay showing the binding efficiency of NeoCoV and PDF-2180-CoV S1-CTD to human and Bat37ACE2-ecto proteins. **e,** The inhibitory activity of soluble ACE2-ecto proteins against NeoCoV infection in 293T-Bat37ACE2. **f,** Dose-dependent competition of NeoCoV infection by Bat37ACE2-ecto proteins in 293T-Bat37ACE2 cells. **g,** The inhibitory effect of NeoCoV, PDF-2180-CoV S1-CTD-hFc and MERS-CoV RBD-hFC proteins on NeoCoV infection in 293T-Bat37ACE2 cells. **h,** Receptor preference of chimeric viruses with S1-CTD or S1-NTD swap mutations in cells expressing the indicated receptors. Mean±SD for **d**, **e**, **g**, and **h**, (n=3).

### Structural basis of ACE2 binding

To unveil the molecular details of the virus-ACE2 binding, we then carried out structural investigations of the Bat37ACE2 in complex with the NeoCoV and PDF-2180-CoV RBD. 3D classification revealed that the NeoCoV-Bat37ACE2 complex primarily adopts a dimeric configuration with two copies of ACE2 bound to two RBDs, whereas only a monomeric conformation was observed in the PDF-2180-CoV-Bat37ACE2 complex (Figs. 3a-b, Extended Data Fig. 10-11). We determined the structures of these two complexes at a resolution of 3.5 Å and 3.8 Å, respectively, and performed local refinement to further improve the densities around the binding interface, enabling reliable analysis of the interaction details (Figs. 3a-b, Extended Data Fig. 12-13 and Table 1-2). Despite existing in different oligomeric states, the structures revealed that both NeoCoV and PDF-2180-CoV recognized the Bat37ACE2 in a very similar way. We used the NeoCoV-Bat37ACE2 structure for detailed analysis (Figs. 3a-b and Extended Data Fig. 14). Like other structures of homologs, the NeoCoV RBD structure comprises a core subdomain located far away from the engaging ACE2 and an external subdomain recognizing the receptor (Fig. 3c and Extended Data Fig. 15). The external subdomain is a strand-enriched structure with four anti-parallel β strands (β6–β9) and exposes a flat four-stranded sheet-tip for ACE2 engagement (Fig. 3c). By contrast, the MERS-CoV RBD recognizes the side surface of the DPP4 β-propeller via its four-stranded sheet-blade (Fig. 3c). The structural basis for the differences in receptor usage can be inferred from two features: i) the local configuration of the four-stranded sheet in the external domain of NeoCoV shows a conformational shift of η3 and β8 disrupting the flat sheet-face for DPP4 binding and ii) relatively longer 6-7 and 8-9 loops observed in MERS-CoV impair their binding in the shallow cavity of bat ACE2 (Fig. 3c and Extended Data Fig. 15).

**Fig. 3.**
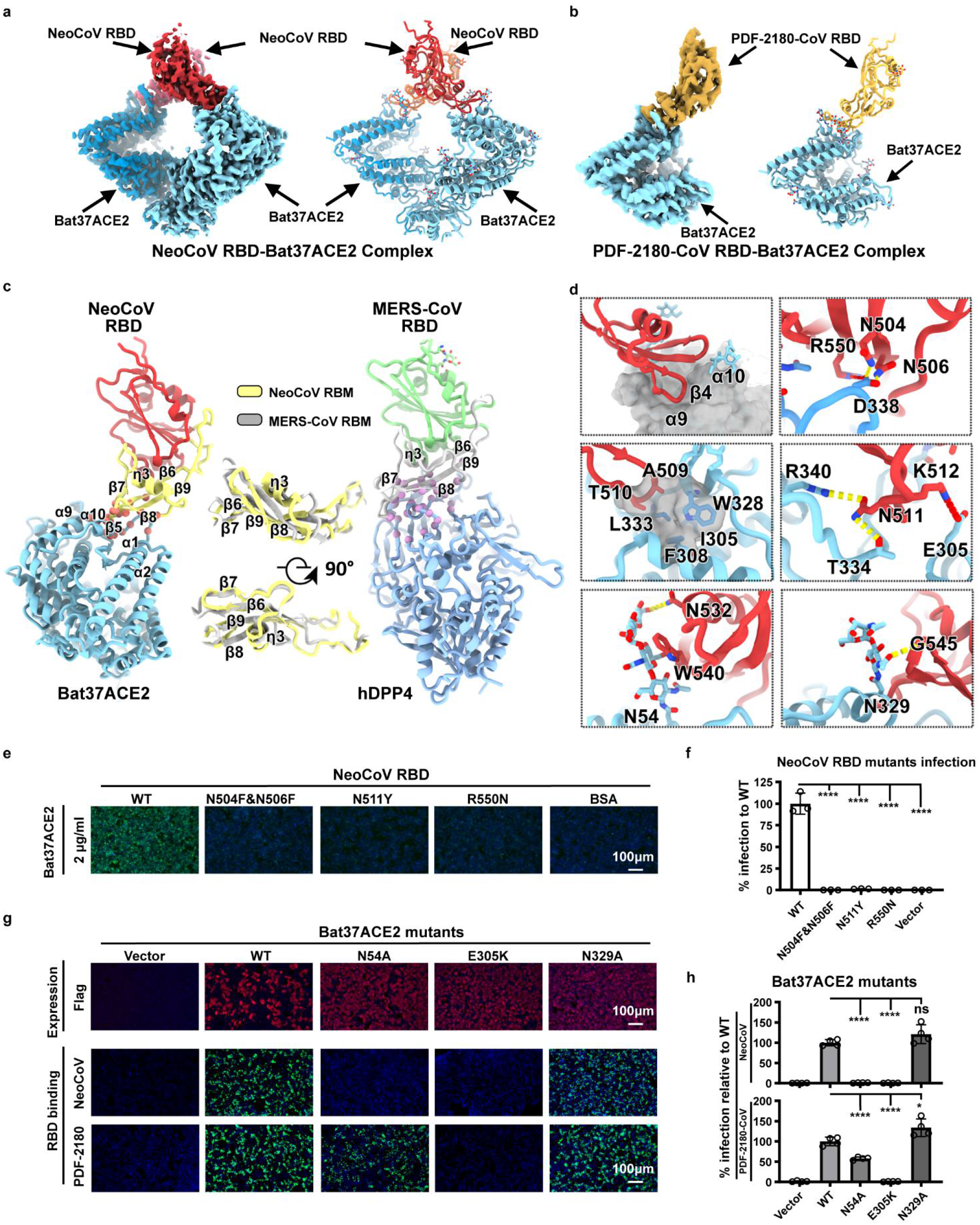
Structure of the NeoCoV RBD-Bat37ACE2 and PDF-2018-CoV RBD-Bat37ACE2 complex. **a**-**b**, Cryo-EM density map and cartoon representation of NeoCoV RBD-Bat37ACE2 complex (**a**) and PDF-2018CoV RBD-Bat37ACE2 complex (**b**). The NeoCoV RBD, PDF-2180-CoV RBD, and Bat37ACE2 were colored by red, yellow, and cyan, respectively. **c**, Structure comparison between NeoCoV RBD-Bat37ACE2 complex (left) and MERS-CoV RBD-hDPP4 complex (right). The NeoCoV RBD, MERS-CoV RBD, NeoCoV RBM, MERS-CoV RBM, Bat37ACE2, and hDPP4 were colored in red, light green, light yellow, gray, cyan, and blue, respectively. **d**, Details of the NeoCOV RBD-Bat37ACE2 complex interface. All structures are shown as ribbon with the key residues shown with sticks. The salt bridges and hydrogen bonds are presented as red and yellow dashed lines, respectively. **e**-**f**, Verification of the critical residues on NeoCoV RBD affecting viral binding (**e**), and entry efficiency (**f**) in 293T-Bat37ACE2 cells. **g**-**h**, Verification of the critical residues on Bat37ACE2 affecting NeoCoV RBD binding (**g**), and viral entry efficiency(**h**). Mean±SD for **f** (n=3) and **h** (n=4).

In the NeoCoV-Bat37ACE2 complex structure, relatively smaller surface areas (498 Å^2^ in NeoCoV RBD and 439 Å^2^ in Bat37ACE2) are buried by the two binding entities compared to their counterparts in the MERS-CoV-DPP4 complex (880 Å^2^ in MERS-CoV RBD and 812 Å^2^ in DPP4; 956 Å^2^ in SARS-CoV-2 RBD and 893 Å^2^ in hACE2). The NeoCoV RBD inserts into an apical depression constructed by α11, α12 helices and a loop connecting α12 and β4 of Bat37ACE2 through its four-stranded sheet tip (Fig. 3d and Extended Data Table. 2). Further examination of the binding interface revealed a group of hydrophilic residues at the site, forming a network of polar-contacts (H-bond and salt-bridge) network and hydrophobic interactions. These polar interactions are predominantly mediated by the residues N504, N506, N511, K512, and R550 from the NeoCoV RBM and residues T53, E305, T334, D338, R340 from Bat37ACE2 (Fig. 3d, Extended Data Table. 2). Notably, the methyl group from residues A509 and T510 of the NeoCoV RBM are partially involved in forming a hydrophobic pocket with residues F308, W328, L333, and I358 from Bat37ACE2 at the interface. A substitution of T510 with F in the PDF-2180-CoV RBM further improves hydrophobic interactions, which is consistent with an increased binding affinity observed for this point mutation (Figs. 3d, Extended Data Table. 2). Apart from protein-protein contacts, the glycans of bat ACE2 at positions N54 and N329 sandwich the strands (β8–β9), forming π-π interactions with W540 and hydrogen bonds with N532, G545, and R550 from the NeoCoV RBD, underpinning virus-receptor associations (Fig. 3d and Extended Data Table. 2).

The critical residues were verified by introducing mutations and testing their effect on receptor binding and viral entry. As expected, mutations N504F/N506F, N511Y, and R550N in the NeoCoV RBD, abolishing the polar-contacts or introducing steric clashes, resulted in a significant reduction of RBD binding and viral entry (Fig.3e-f). Similarly, E305K mutation in Bat37ACE2 eliminating the salt-bridge also significantly impaired the receptor function. Moreover, the loss of function effect of mutation N54A on Bat37ACE2 abolishing the N-glycosylation at residue 54 confirmed the importance of the particular protein-glycan interaction in viral-receptor recognition. In comparison, N329A abolishing the N-glycosylation at site N329, located far away from the binding interface, had no significant effect on receptor function (Fig.3g-h).

### Evaluation of zoonotic potential

A major concern is whether NeoCoV and PDF-2180-CoV could jump the species barrier and infect humans. As mentioned above, NeoCoV and PDF-2180-CoV cannot efficiently interact with human ACE2. Here we first examined the molecular determinants restricting hACE2 from supporting the entry of these viruses. By comparing the binding interface of the other three hACE2-using coronaviruses, we found that the SARS-CoV, SARS-COV-2, and NL63 share similar interaction regions that barely overlapped with the region engaged by NeoCoV (Fig. 4a). Analysis of the overlapped binding interfaces reveals a commonly used hot spot around residues 329-330 (Fig.4b). Through sequences alignment and structural analysis of hACE2 and Bat37ACE2, we predicted that the inefficient use of the hACE2 for entry by the viruses could be attributed to incompatible residues located around the binding interfaces, especially the difference in sequences between residues 337-342 (Fig.4c). We replaced these residues of hACE2 with those from the Bat37ACE2 counterparts to test this hypothesis (Fig.4c-d). The substitution led to an approximately 15-fold and 30-fold increase in entry efficiency of NeoCoV and PDF-2180-CoV, respectively, confirming that this region is critical for the determination of the host range. Further fine-grained dissection revealed that N338 is the most crucial residue in restricting human receptor usage (Fig. 4e-g).

**Fig. 4.**
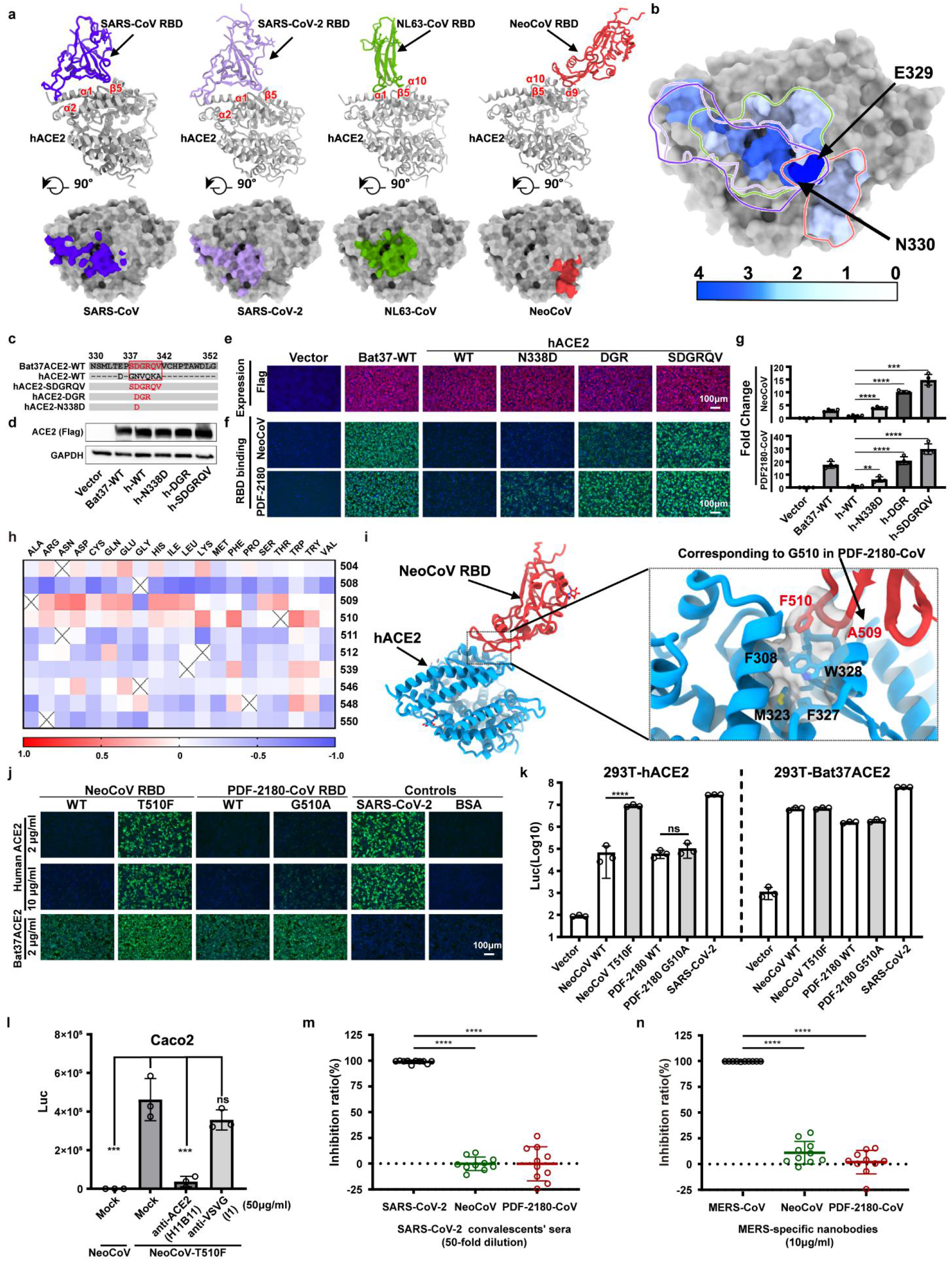
Molecular determinants affecting hACE2 recognition by the viruses. **a**, Binding modes of ACE2-adapted coronaviruses. The SARS-CoV RBD, SARS-CoV-2 RBD, NL63-CoV RBD, and NeoCoV RBD were colored in purple, light purple, green, and red, respectively. **b**, A common virus-binding hot spot on ACE2 for the four viruses. Per residue frequency recognized by the coronavirus RBDs were calculated and shown. **c**, Schematic illustration of the hACE2 swap mutants with Bat37ACE2 counterparts. **d**-**e**, The expression level of the hACE2 mutants by Western blot (**d**) and immunofluorescence (**e**). **f**-**g**, Receptor function of hACE2 mutants evaluated by virus RBD binding assay (**f**) and pseudovirus entry assay (**g**). **h**, Molecular dynamics (MD) analysis of the effect of critical residue variations on the interaction between NeoCoV and Bat37ACE2 by mCSM-PPI2. **i**, Structure of NeoCoV RBD-hACE2 complex modeling by superposition in COOT. The NeoCoV RBD and hACE2 were colored in red and sky blue, respectively. Details of the NeoCoV RBD key mutation T510F was shown. All structures are presented as ribbon with the key residues shown with sticks. **j-k,** The effect of NeoCoV and PDF-2180-CoV RBM mutations on hACE2 fitness as demonstrated by binding **(j**) and entry efficiency (**k**) on 293T-hACE2 and 293T-Bat37ACE2 cells. **l,** hACE2 dependent entry of NeoCoV-T510F in Caco2 cells in the presence of 50μg/ml of Anti-ACE2 (H11B11) or Anti-VSVG (I1). **m,** Neutralizing activity of SARS-CoV-2 vaccinated sera against the infection by SARS-CoV-2, NeoCoV, and PDF-2180-CoV. **n,** Neutralizing activity of MERS-RBD targeting nanobodies against the infections by MERS-CoV, NeoCoV, and PDF-2180-CoV. Mean± SD for **g**,**k-n**; **g**(n=4),**k**-**l** (n=3), g**m**-**n** (n=10).

We further assessed the zoonotic potential of NeoCoV and PDF-2180-CoV by identifying the molecular determinants of viral RBM, which might allow cross-species transmission through engaging hACE2. After meticulously examining the critical residues based on the complex structures and computational prediction tool mCSM-PPI2^33^(Fig. 4h, Extended Data Table. 4), we predicted increasing hydrophobicity around the residue T510 of NeoCoV might enhance the virus-receptor interaction on hACE2 (Fig. 4 i). Interestingly, the PDF-2180-CoV already has an F511 (corresponding to site 510 of NeoCoV), which is consistent with its slightly higher affinity to human ACE2 (Extended Data Fig.16). As expected, T510F substitutions in NeoCoV remarkably increased its binding affinity with hACE2 (KD=16.9 nM) and a significant gain of infectivity in 293T-hACE2 cells (Fig. 4 j-k, Extended Data Fig.17-18). However, PDF-2180-CoV showed much lower efficiency in using hACE2 than NeoCoV-T510F, indicating other unfavorable residues are restricting its efficient interaction with hACE2. Indeed, a G to A (corresponding to A509 in NeoCoV) mutation in site 510, increasing the local hydrophobicity, partially restored its affinity to hACE2 (Fig.4 j-k). In addition, the NeoCoV-T510F can enter the human colon cell line Caco-2 with much higher efficiency than wild-type NeoCoV. It enters the Caco-2 exclusively through ACE2 as the infection can be neutralized by an ACE2-targeting neutralizing antibody H11B11^34^ (Fig. 4l). Humoral immunity triggered by prior infection or vaccination of other coronaviruses might be inadequate to protect humans from NeoCoV and PDF-2180-CoV infections because neither SARS-CoV-2 anti-sera nor ten tested anti-MERS-CoV nanobodies can cross-inhibit the infection caused by these two viruses^35^. (Fig. 4m-n).

## Discussion

The lack of knowledge of the receptors of bat coronaviruses has greatly limited our understanding of these high-risk pathogens. Our study provided evidence that the relatives of potential MERS-CoV ancestors like NeoCoV and PDF-2180-CoV engage bat ACE2 for efficient cellular entry. However, HKU5-CoV and EriCoV seem not to use bat DPP4 or hedgehog ACE2 for entry, highlighting the complexity of coronaviruses receptor utilization. It was unexpected that NeoCoV and PDF-2180-CoV use ACE2 rather than DPP4 as their entry receptors since their RBD core structures resemble MERS-CoV more than other ACE2-using viruses (Fig. 4a, Extended Data Fig. 15).

Different receptor usage can affect the transmission rate of the viruses. Although it remains unclear whether ACE2 usage out-weight DPP4 usage for more efficient transmission, MERS-CoV appears to have lower transmissibility with an estimated R0 around 0.69. Comparatively, the ACE2 usage has been approved able to achieve high transmissibility. The SARS-CoV-2 estimated R0 is around 2.5 for the original stain, 5.08 for the delta variant, and even higher for the omicron variant^36–38^. This unexpected ACE2 usage of these MERS-CoV close relatives highlights a latent biosafety risk, considering a combination of two potentially damaging features of high fatality observed for MERS-CoV and the high transmission rate noted for SARS-CoV-2. Furthermore, our studies show that the current COVID-19 vaccinations are inadequate to protect humans from any eventuality of the infections caused by these viruses.

Many sarbecoviruses, alpha-CoV NL63, and a group of merbecoviruses reported in this study share ACE2 for cellular entry. Our structural analysis indicates NeoCoV and PDF-2180-CoV bind to an apical side surface of ACE2, which is different from the surface engaged by other ACE2-using coronaviruses (Fig.4a). The interaction is featured by inter-molecular protein-glycan bonds formed by the glycosylation at N54, which is not found in RBD-receptor interactions of other coronaviruses. The different interaction modes of the three ACE2-using coronaviruses indicate a history of multiple independent receptor acquisition events during evolution^22^. The evolutionary advantage of ACE2 usage in different CoVs remains enigmatic.

Our results support the previous hypothesis that the origin of MERS-CoV might be a result of an intra-spike recombination event between a NeoCoV like virus and a DPP4-using virus^26^. RNA recombination can occur during the co-infection of different coronaviruses, giving rise to a new virus with different receptor usage and host tropisms^39, 40^. It remains unclear whether the event took place in bats or camels, and where the host switching happened. Although bat merbecoviruses are geographically widespread, the two known ACE2-using merbecoviruses are inhabited in Africa. Moreover, most camels in the Arabian Peninsula showing serological evidence of previous MERS-CoV infection are imported from the Greater Horn of Africa with several Neoromicia species^41^. Considering both viruses are inefficient in infecting human cells in their current form, the acquisition of the hDPP4 binding domain would be a critical event driving the emergence of MERS-CoV. Further studies will be necessary to obtain more evidence about the origin of MERS-CoV.

The host range determinants on ACE2 are barriers for cross-species transmission of these viruses. Our results show NeoCoV and PDF-2180-CoV favor ACE2 from bats of the *Yangochiroptera* group, especially vesper bats (*Vespertilionidae*), where their host belongs to, but not ACE2 orthologs from bats of the *Yinpterochiroptera* group. Interestingly, most merbecoviruses were found in species belonging to the *Vespertilionidae* group, a highly diverse and widely distributed family^9^. Although the two viruses could not use hACE2 efficiently, our study also reveals that single residue substitution increasing local hydrophobicity around site 510 could enhance their affinity for hACE2 and enable them to infect human cells expressing ACE2. Considering the extensive mutations in the RBD regions of the SARS-CoV-2 variants, especially the heavily mutated omicron variant, these viruses may hold a latent potential to infect humans through further adaptation via antigenic drift^42, 43^. It is also very likely that their relatives with human emergence potential are circulating somewhere in nature.

Overall, we identified ACE2 as a long-sought functional receptor of the potential MERS-CoV ancestors in bats, facilitating the in-depth research of these important viruses with zoonotic emergence risks. Our study adds to the knowledge about the complex receptor usage of coronaviruses, highlighting the importance of surveillance and research on these viruses to prepare for potential outbreaks in the future.

## Supplementary Information

### Methods

#### Receptor and virus sequences

The acquisition of sequences of 46 bat ACE2 and hACE were described in our previous study^31^. The five bat DPP4 and hDPP4 sequences were directly retrieved from the GenBank database (human DPP4, NM_001935.3; Bat37, Pipistrellus pipistrellus, KC249974.1) or extracted from whole genome sequence assemblies of the bat species retrieved from the GenBank database (Bat25, Sturnira hondurensis, GCA_014824575.2; Bat29, Mormoops blainvillei, GCA_004026545.1; Bat36, Aeorestes cinereus, GCA_011751065.1; Bat40, Antrozous pallidus, GCA_007922775.1). The whole genome sequences of different coronaviruses were retrieved from the GenBank database. The accession numbers are as follows: MERS-CoV (JX869059.2), Camel MERS-CoV KFU-HKU 19Dam (KJ650296.1), HKU4 (NC_009019.1), HKU5 (NC_009020.1), ErinaceusCoV/HKU31 strain F6 (MK907286.1), NeoCoV (KC869678.4), PDF-2180-CoV (NC_034440.1), ErinaceusCoV/2012-174 (NC_039207.1), BtVs-BetaCoV/SC2013 (KJ473821.1), BatCoV/H.savii/Italy (MG596802.1), BatCoV HKU25 (KX442564.1), BatCoV ZC45(MG772933.1) and SARS-CoV-2 (NC_045512.2), NL63 (JX504050.1) 229E (MT797634.1).

All gene sequences used in this study were commercially synthesized by Genewiz. The sources, accession numbers, and sequences of the receptors and viruses were summarized in Supplementary Table 1.

#### SARS-CoV-2 anti-sera collection

All the vaccinated sera were collected from volunteers at about 21 days post the third dose of the WHO-approved inactivated SARS-COV-2 vaccine (CorovaVac, Sinovac, China). All volunteers were provided informed written consent forms, and the whole study was conducted following the requirements of Good Clinical Practice of China.

#### Bioinformatic analysis

Protein sequence alignment was performed using the MUSCLE algorithm by MEGA-X software (version 10.1.8). For phylogenetic analysis, nucleotide or protein sequences of the viruses were first aligned using the Clustal W and the MUSCLE algorithm, respectively. Then, the phylogenetic trees were generated using the maximal likelihood method in MEGA-X (1000 Bootstraps). The model and the other parameters used for phylogenetic analysis were applied following the recommendations after finding best DNA/Protein Models by the software. The nucleotide similarity of coronaviruses was analyzed by SimPlot software (version 3.5.1) with a slide windows size of 1000 nucleotides and a step size of 100 nucleotides using gap-stripped alignments and the Kimura (2-parameter) distance model.

#### Plasmids

Human codon-optimized sequences of various ACE2 or DPP4 orthologs and their mutants were cloned into a lentiviral transfer vector (pLVX-IRES-puro) with a C-terminal 3×Flag tag (DYKDHD-G-DYKDHD-I-DYKDDDDK). The DNA sequences of human codon-optimized NeoCoV S protein (AGY29650.2), PDF-2180-CoV S protein (YP_009361857.1), HKU4-CoV S protein (YP_001039953.1), HKU5-CoV S protein (YP_001039962.1), HKU31 S protein (QGA70692.1), SARS-CoV-2 (YP_009724390.1), and MERS-CoV S protein (YP_009047204.1) were cloned into the pCAGGS vector with a C-terminal 13-15-amino acids deletion (corresponding to 18 amino-acids in SARS-CoV-2) or replacement by an HA tag (YPYDVPDYA) for higher VSV pseudotyping efficiency^44^. The plasmids expressing coronavirus RBD-IgG-hFc fusion proteins were generated by inserting the coding sequences of NeoCoV RBD (aa380-585), PDF-2180-CoV RBD (aa381-586), HKU4-CoV (aa382-593), HKU5-CoV RBD (aa385-586), HKU31-CoV RBD (aa366-575), SARS-CoV-2 RBD (aa331-524) and MERS-CoV RBD (aa377-588) into the pCAGGS vector with an N-terminal CD5 secretion leading sequence (MPMGSLQPLATLYLLGMLVASVL). The plasmids expressing soluble bat ACE2 and DPP4 proteins were constructed by inserting the ectodomain coding sequences into the pCAGGS vector with N-terminal CD5 leader sequence and C-terminal twin-strep tag and 3×Flag tag tandem sequences (WSHPQFEKGGGSGGGSGGSAWSHPQFEK-GGGRS-DYKDHDGDYKDHDIDYKDDDDK).

Virus spike proteins or receptor-related mutants or chimeras were generated by overlapping PCR. For Dual split protein (DSP) based cell-cell fusion assay, the dual reporter split proteins were expressed by pLVX-IRES-puro vector expressing the RLucaa1-155-GFP1-7(aa1-157) and GFP8-11(aa158-231)-RLuc-aa156-311 plasmids, which were constructed in the lab based on a previously study^32, 45^. The plasmids expressing the codon-optimized anti-ACE2 antibody (H11B11; GenBank accession codes MZ514137 and MZ514138) were constructed by inserting the heavy-chain and light-chain coding sequences into the pCAGGS vector with N-terminal CD5 leader sequences, respectively^34^. For anti-MERS-CoV nanobody-hFc fusion proteins, nanobody coding sequences were synthesized and cloned into the pCAGGS vector with N-terminal CD5 leader sequences and C-terminal hFc tags ^35^.

#### Protein expression and purification

The RBD-hFc (S1-CTD-hFc) fusion proteins of SARS-CoV-2, MERS-CoV, HKU4-CoV, HKU5-CoV, HKU31-CoV, NeoCoV, and PDF-2180-CoV, and the soluble ACE2 proteins of human, Bat25, Bat29, Bat36, Bat37, Bat38, and Bat40 were expressed by 293T by transfecting the corresponding plasmids by GeneTwin reagent (Biomed, TG101-01) following the manufacturers’ instructions. Four hrs post-transfection, the culture medium of the transfected cells was replenished by SMM 293-TII Expression Medium (Sino Biological, M293TII). The supernatant of the culture medium containing the proteins was collected every 2-3 days. The recombinant RBD-hFc proteins were captured by Pierce Protein A/G Plus Agarose (Thermo Scientific, 20424), washed by wash buffer W (100 mM Tris/HCl, pH 8.0, 150 mM NaCl, 1mM EDTA), eluted with pH 3.0 Glycine buffer (100mM in H2O), and then immediately balanced by UltraPure 1M Tris-HCI, pH 8.0 (15568025, Thermo Scientific). The twin-strep tag containing proteins were captured by Strep-Tactin XT 4Flow high capacity resin (IBA, 2-5030-002), washed by buffer W, and eluted with buffer BXT (100 mM Tris/HCl, pH 8.0, 150 mM NaCl, 1mM EDTA, 50mM biotin). The eluted proteins can be concentrated and buffer-changed to PBS through ultra-filtration. Protein concentrations were determined by Omni-Easy Instant BCA Protein Assay Kit (Epizyme, ZJ102). The purified proteins were aliquoted and stored at -80℃. For Cryo-EM analysis, NeoCoV RBD (aa380-588), PDF-2018-CoV RBD (381-589), and Bat37ACE2 (aa21-730) were synthesized and subcloned into the vector pCAGGS with a C-terminal twin-strep tag. Briefly, these proteins were expressed by transient transfection of 500 ml HEK Expi 293F cells (Gibco, Thermo Fisher, A14527) using Polyethylenimine Max Mw 40,000 (polysciences). The resulting protein samples were further purified by size-exclusion chromatography using a Superdex 75 10/300 Increase column (GE Healthcare) or a Superdex 200 10/300 Increase column (GE Healthcare) in 20mM HEPES, 100 mM NaCl, pH 7.5. For RBD-receptor complex (NeoCoV RBD-Bat37ACE2 / PDF-2180-CoV RBD-Bat37ACE2), NeoCoV RBD or PDF-2180-CoV RBD was mixed with Bat37ACE2 at the ratio of 1.2 :1, incubated for 30 mins on ice. The mixture was then subjected to gel filtration chromatography. Fractions containing the complex were collected and concentrated to 2 mg/ml.

#### Cell culture

293T (CRL-3216), VERO E6 cells (CRL-1586), A549 (CCL-185), BHK-21 (CCL-10), and Huh-7 (PTA-4583), Caco2 (HTB-37) and the epithelial bat cell line Tb 1 Lu (CCL-88) were purchased from American Type Culture Collection (ATCC) and cultured in Dulbecco’s Modified Eagle Medium, (DMEM, Monad, China) supplemented with 10% fetal bovine serum (FBS), 2.0 mM of L-glutamine, 110 mg/L of sodium pyruvate and 4.5 g/L of D-glucose. An I1-Hybridoma (CRL-2700) cell line secreting a neutralizing mouse monoclonal antibody against the VSV glycoprotein (VSVG) was cultured in Minimum Essential Medium with Earles’s balances salts and 2.0mM of L-glutamine (Gibico) and 10% FBS. All cells were cultures at 37℃ in 5% CO2 with the regular passage of every 2-3 days. 293T stable cell lines overexpressing ACE2 or DPP4 orthologs were maintained in a growth medium supplemented with 1 μg/ml of puromycin.

#### Stable cell line generation

Stable cell lines overexpressing ACE2 or DPP4 orthologs were generated by lentivirus transduction and antibiotic selection. Specifically, the lentivirus carrying the target gene was produced by cotransfection of lentiviral transfer vector (pLVX-EF1a-Puro, Genewiz) and packaging plasmids pMD2G (Addgene, plasmid no.12259) and psPAX2 (Addgene, plasmid no.12260) into 293T cells through Lip2000 Transfection Reagent (Biosharp, BL623B). The lentivirus-containing supernatant was collected and pooled at 24 and 48 hrs post-transfection. 293T cells were transduced by the lentivirus after 16 hrs in the presence of 8 μg/ml polybrene. Stable cells were selected and maintained in the growth medium with puromycin (1-2 μg/ml). Cells selected for at least ten days were considered stable cell lines and used in different experiments.

#### Cryo-EM sample preparation and data collection

For Cryo-EM sample preparation, the NeoCoV RBD-Bat37ACE2 or PDF-2018-CoV RBD-Bat37ACE2 complex was diluted to 0.5 mg/ml. Holy-carbon gold grid (Cflat R1.2/1.3 mesh 300) were freshly glow-discharged with a Solarus 950 plasma cleaner (Gatan) for 30s. A 3 μL aliquot of the mixture complex was transferred onto the grids, blotted with filter paper at 16℃ and 100% humidity, and plunged into the ethane using a Vitrobot Mark IV (FEI). For these complexes, micrographs were collected at 300 kV using a Titan Krios microscope (Thermo Fisher), equipped with a K2 detector (Gatan, Pleasanton, CA), using SerialEM automated data collection software. Movies (32 frames, each 0.2 s, total dose 60^e−Å−2^) were recorded at a final pixel size of 0.82 Å with a defocus of between -1.2 and -2.0 μm.

#### Image processing

For NeoCoV RBD-Bat37ACE2 complex, a total of 4,234 micrographs were recorded. For PDF-2018-CoV RBD-Bat37ACE2 complex, a total of 3,298 micrographs were recorded. Both data sets were similarly processed. Firstly, the raw data were processed by MotionCor2, which were aligned and averaged into motion-corrected summed images. Then, the defocus value for each micrograph was determined using Gctf. Next, particles were picked and extracted for two-dimensional alignment. The well-defined partial particles were selected for initial model reconstruction in Relion^46^. The initial model was used as a reference for three-dimensional classification. After the refinement and post-processing, the overall resolution of PDF-2018-CoV RBD-Bat37ACE2 complex was up to 3.8Å based on the gold-standard Fourier shell correlation (threshold = 0.143) ^47^. For the NeoCoV RBD-Bat37ACE2 complex, the C2 symmetry was expanded before the 3D refinement. Finally, the resolution of the NeoCoV RBD-Bat37ACE2 complex was up to 3.5Å. The quality of the local resolution was evaluated by ResMap^48^.

#### Model building and refinement

The NeoCoV RBD-Bat37ACE2 complex structures were manually built into the refined maps in COOT^47^. The atomic models were further refined by positional and B-factor refinement in real space using Phenix^48^. For the PDF-2018-CoV RBD-Bat37ACE2 complex model building, the refinement NeoCoV RBD-Bat37ACE2 complex structures were manually docked into the refined maps using UCSF Chimera and further corrected manually by real-space refinement in COOT. As the same, the atomic models were further refined by using Phenix. Validation of the final model was performed with Molprobity^48^. The data sets and refinement statistics are shown in Extended Data table 1.

#### Immunofluorescence assay

The expression levels of ACE2 or DPP4 receptors were evaluated by immunofluorescence assay detecting the C-terminal 3×FLAG-tags. The cells expressing the receptors were seeded in the 96-well plate (poly-lysine pretreated plates for 293T based cells) at a cell density of 1∼5×10^5^/ml (100 μl per well) and cultured for 24 hrs. Cells were fixed with 100% methanol at room temperature for 10 mins, and then incubated with a mouse monoclonal antibody (M2) targeting the FLAG-tag (Sigma-Aldrich, F1804) diluted in 1% BSA/PBS at 37℃ for 1 hour. After one wash with PBS, cells were incubated with 2 μg/ml of the Alexa Fluor 594-conjugated goat anti-mouse IgG (Thermo Fisher Scientific, A32742) diluted in 1% BSA/PBS at room temperature for 1 hour. The nucleus was stained blue with Hoechst 33342 (1:5,000 dilution in PBS). Images were captured with a fluorescence microscope (Mshot, MI52-N).

#### Pseudovirus production and titration

Coronavirus spike protein pseudotyped viruses (CoV-psV) were packaged according to a previously described protocol based on a replication-deficient VSV-based rhabdoviral pseudotyping system (VSV-dG). The VSV-G glycoprotein-deficient VSV coexpressing GFP and firefly luciferase (VSV-dG-GFP-fLuc) was rescued by a reverse genetics system in the lab and helper plasmids from Karafast. For CoV-psV production, 293T or Vero E6 cells were transfected with the plasmids overexpressing the coronavirus spike proteins through the Lip2000 Transfection Reagent (Biosharp, BL623B). After 36 hrs, the transfected cells were transduced with VSV-dG-GFP-fLuc diluted in DMEM for 4 hrs at 37℃ with a 50 % tissue culture infectious dose (TCID50) of 1×10^6^ TCID50/ml. Transduced cells were washed once with DMEM and then incubated with culture medium and I1-hybridoma-cultured supernatant (1:10 dilution) containing VSV neutralizing antibody to eliminate the infectivity of the residual input viruses. The CoV-psV-containing supernatants were collected at 24 hrs after the medium change, clarified at 4,000 rpm for 5 mins, aliquoted, and stored at -80℃. The TCID50 of pseudovirus was determined by a serial-dilution based infection assay on 293T-bat40ACE2 cells for NeoCoV and PDF-2180-CoV or 293T-hDpp4 cells for MERS-CoV and HKU4-CoV. The TCID50 was calculated according to the Reed-Muench method ^49, 50^. The relative luminescence unit (RLU) value ≥ 1000 is considered positive. The viral titer (genome equivalents) of HKU5-COV and HKU31-CoV without an ideal infection system was determined by quantitative PCR with reverse transcription (RT–qPCR). The RNA copies in the virus-containing supernatant were detected using primers in the VSV-L gene sequences (VSV-L-F: 5’-TTCCGAGTTATGGGCCAGTT-3’; VSVL-R: 5’-TTTGCCGTAGACCTTCCAGT-3’).

#### Pseudovirus entry assay

Cells for infection were trypsinized and incubated with different pseudoviruses (1×10^5^ TCID50/well, or same genome equivalent) in a 96-well plate (5×10^4^ /well) to allow attachment and viral entry simultaneously. For TPCK-trypsin treatment for infection boosting, NeoCoV and PDF-2180-CoV pseudovirus in serum-free DMEM were incubated with 100 μg/ml TPCK-treated trypsin (Sigma-Aldrich, T8802) for 10 mins at 25℃, and then treated with 100 μg/ml soybean trypsin inhibitor (Sigma-Aldrich, T6414) in DMEM+10% FBS to stop the proteolysis. At 16 hours post-infection (hpi), GFP images of infected cells were acquired with a fluorescence microscope (Mshot, MI52-N), and intracellular luciferase activity was determined by a Bright-Glo Luciferase Assay Kit (Promega, E2620) and measured with a SpectraMax iD3 Multi-well Luminometer (Molecular Devices) or a GloMax 20/20 Luminometer (Promega).

#### Pseudovirus neutralization Assay

For antibody neutralization assays, the viruses (2×10^5^ TCID50/well) were incubated with the sera (50-fold diluted) or 10 μg/ml MERS-specific nanobodies at 37℃ for 30 mins, and then mixed with trypsinized BHK-21-Bat37ACE2 cells with the density of 2×10^4^/well. After 16 hrs, the medium of the infected cells was removed, and the cells were lysed with 1×Bright-Glo Luciferase Assay reagent (Promega) for chemiluminescence detection with a SpectraMax iD3 Multi-well Luminometer (Molecular devices).

#### Western blot

After one wash with PBS, the cells were lysed by 2% TritonX-100/PBS containing 1mM fresh prepared PMSF (Beyotime, ST506) on ice for 10 mins. Then cell lysate was clarified by 12,000 rpm centrifugation at 4℃ for 5 mins, mixed with SDS loading buffer, and then incubated at 95 °C for 5 mins. After SDS-PAGE electrophoresis and PVDF membrane transfer, the membrane was blocked with 5% skim milk/PBST at room temperature for one hour, incubated with primary antibodies against Flag (Sigma, F1804), HA (MBL, M180-3), or glyceraldehyde-3-phosphate dehydrogenase (GAPDH) (AntGene, ANT011) at 1:10000 dilution in 1% milk/PBS overnight on a shaker at 4℃. After extensive wash, the membrane was incubated with the Horseradish peroxidase (HRP)-conjugated secondary antibody AffiniPure Goat Anti-Mouse IgG (H+L) (Jackson Immuno Reseach, 115-035-003) in 1% skim milk in PBST, and incubated for one hour. The blots were visualized using Omni-ECL Femto Light Chemiluminescence Kit (EpiZyme, SQ201) by ChemiDoc MP (Bio-Rad).

#### Coronavirus RBD-hFc live-cell binding assay

Recombinant coronavirus RBD-hFc proteins (1-16 μg/ml) were diluted in DMEM and then incubated with the cells for one hour at 37℃. Cells were washed once with DMEM and then incubated with 2 μg/ml of Alexa Fluor 488-conjugated goat anti-human IgG (Thermo Fisher Scientific; A11013) diluted in Hanks’ Balanced Salt Solution (HBSS) with 1% BSA for 1 hour at 37 ℃. Cells were washed twice with PBS and incubated with Hoechst 33342 (1:5,000 dilution in HBSS) for nucleus staining. Images were captured with a fluorescence microscope (MI52-N). For flow cytometry analysis, cells were detached by 5mM of EDTA/PBS and analyzed with a CytoFLEX Flow Cytometer (Beckman).

#### Biolayer interferometry (BLI) binding assay

The protein binding affinities were determined by BLI assays performed on an Octet RED96 instrument (Molecular Devices). Briefly, 20 μg/mL Human Fc-tagged RBD-hFc recombinant proteins were loaded onto a Protein A (ProA) biosensors (ForteBio, 18-5010) for 30s. The loaded biosensors were then dipped into the kinetic buffer (PBST) for 90s to wash out unbound RBD-hFc proteins. Subsequently, the biosensors were dipped into the kinetic buffer containing soluble ACE2 with concentrations ranging from 0 to 500 nM for 120s to record association kinetic and then dipped into kinetics buffer for 300s to record dissociation kinetics. Kinetic buffer without ACE2 was used to define the background. The corresponding binding affinity (KD) was calculated with Octet Data Analysis software 12.2.0.20 using curve-fitting kinetic analysis or steady-state analysis with global fitting.

#### Enzyme-linked immunosorbent assay (ELISA)

To evaluate the binding between viral RBD and the ACE2 *in vitro*, 96 well Immuno-plate were coated with ACE2 soluble proteins at 27 μg/ml in BSA/PBS (100 μl/well) overnight at 4℃. After three wash by PBS containing 0.1% Tween-20 (PBST), the wells were blocked by 3% skim milk/PBS at 37℃ for 2 hrs. Next, varying concentrations of RBD-hFc proteins (1-9 μg/ml) diluted in 3% milk/PBST were added to the wells and incubated for one hour at 37℃. After extensive wash, the wells were incubated with 1:2000 diluted HRP-conjugated goat anti-human Fc antibody (Sigma, T8802) in PBS for one hour. Finally, the substrate solution (Solarbio, PR1210) was added to the plates, and the absorbance at 450nm was measured with a SpectraMax iD3 Multi-well Luminometer (Molecular Devices).

### Cell-cell fusion assay

Cell-cell fusion assay based on Dual Split proteins (DSP) was conducted on BHK-21 cells stably expressing different receptors^32^. The cells were separately transfected with Spike and RLuc_aa1-155_-GFP_1-10(aa1-157)_ expressing plasmids, and Spike and GFP_11(aa158-231)_ RLuc-C_aa156-311_ expressing plasmids, respectively. At 12 hrs after transfection, the cells were trypsinized and mixed into a 96-well plate at 8×10^4^/well. At 26 hrs post-transfection, cells were washed by DMEM once and then incubated with DMEM with or without 12.5 μg/ml TPCK-trypsin for 25 mins at RT. Five hrs after treatment, the nucleus was stained blue with Hoechst 33342 (1:5,000 dilution in HBSS) for 30min at 37℃. GFP images were then captured with a fluorescence microscope (MI52-N; Mshot). For live-cell luciferase assay, the EnduRen live cell substrate (Promega, E6481) was added to the cells (a final concentration of 30 μM in DMEM) for at least 1 hour before detection by a GloMax 20/20 Luminometer (Promega).

### Statistical Analysis

Most experiments were repeated 2∼5 times with 3-4 biological repeats, each yielding similar results. Data are presented as MEAN±SD or MEAN±SEM as specified in the figure legends. All statistical analyses were conducted using GraphPad Prism 8. Differences between independent samples were evaluated by unpaired two-tailed t-tests; Differences between two related samples were evaluated by paired two-tailed t-tests. P<0.05 was considered significant. * p<0.05, ** p <0.01, *** p <0.005, and **** p <0.001.

## Acknowledgements

We thank Dr. Xiaojun Huang, Dr. Xujing Li, and Dr. Lihong Chen for cryo-EM data collection at the Center for Biological Imaging (CBI) in Institution of Biophysics, CAS. We thank Dr. Yuanyuan Chen, Zhenwei Yang, Bingxue Zhou, and Xianjin Ou for technical support on BLI and SPR. We thank professor Qihui Wang (Institute of Microbiology Chinese Academy of Sciences) for her generous gift of Hedgehog ACE2 plasmid. We thank Ye Fu for his recommendations about manuscript writing. We thank Beijing Taikang Yicai Foundation for their support to this work. This study was supported by China NSFC projects (32070160); Fundamental Research Funds for the Central Universities (2042020kf0024); Special Fund for COVID-19 Research of Wuhan University; The Strategic Priority Research Program (XDB29010000, XDB37030000), CAS (YSBR-010); National Key Research and Development Program (2020YFA0707500, 2018YFA0900801); Xiangxi Wang was supported by Ten Thousand Talent Program and the NSFS Innovative Research Group (No. 81921005).

## Author contributions

H.Y. and X.X.W. conceived and designed the study. Q.X., L.C., C.B.M., C.L., J.Y.S., P.L., and F.T. performed the experiments. Q.X, L.C, C.B.M, C.L, C.F.Z., H.Y, and X.X.W analyzed the data. H.Y., X.X.W., Q.X, L.C, C.B.M, and C.L interpreted the results. H.Y and X.X.W wrote the initial drafts of the manuscript. H.Y., X.X.W., H.Y., X.X.W., L.C., and Q.X. revised the manuscript. C.B.M, C.L., P. L., M.X.G., C.L.W, L.L.S, F.T. M.L.H, J.L., C.S., Y.C., H.B.Z., and K.L. commented on the manuscript.

## Data availability

The cryo-EM maps have been deposited at the Electron Microscopy Data Bank (www.ebi.ac.uk/emdb) and are available under accession numbers: EMD-32686 (NeoCoV RBD-Bat37ACE2 complex) and EMD-32693 (PDF-2180-CoV RBD-Bat37ACE2 complex). Atomic models corresponding to EMD-32686, EMD-32693 have been deposited in the Protein Data Bank (www.rcsb.org) and are available under accession numbers, PDB ID 7WPO, PDB ID 7WPZ, respectively. The authors declare that all other data supporting the findings of this study are available with the paper and its supplementary information files.

## Additional Information

Supplementary Information is available for this paper.

Correspondence and requests for materials should be addressed to H.Y.

---

**Extended Data Figure 1.**
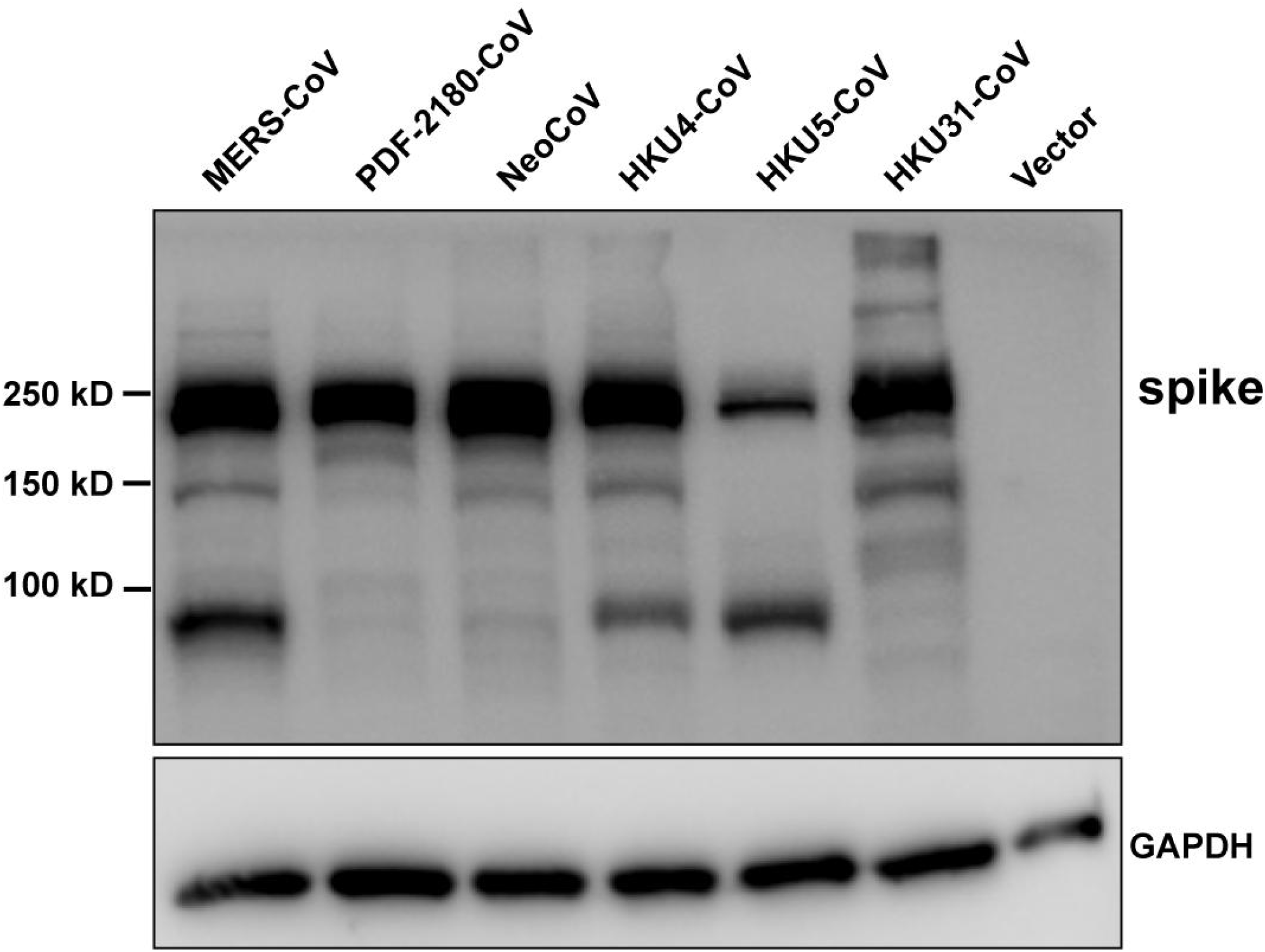
Expression level of coronaviruses spike proteins used for pseudotyping.

**Extended Data Figure 2.**
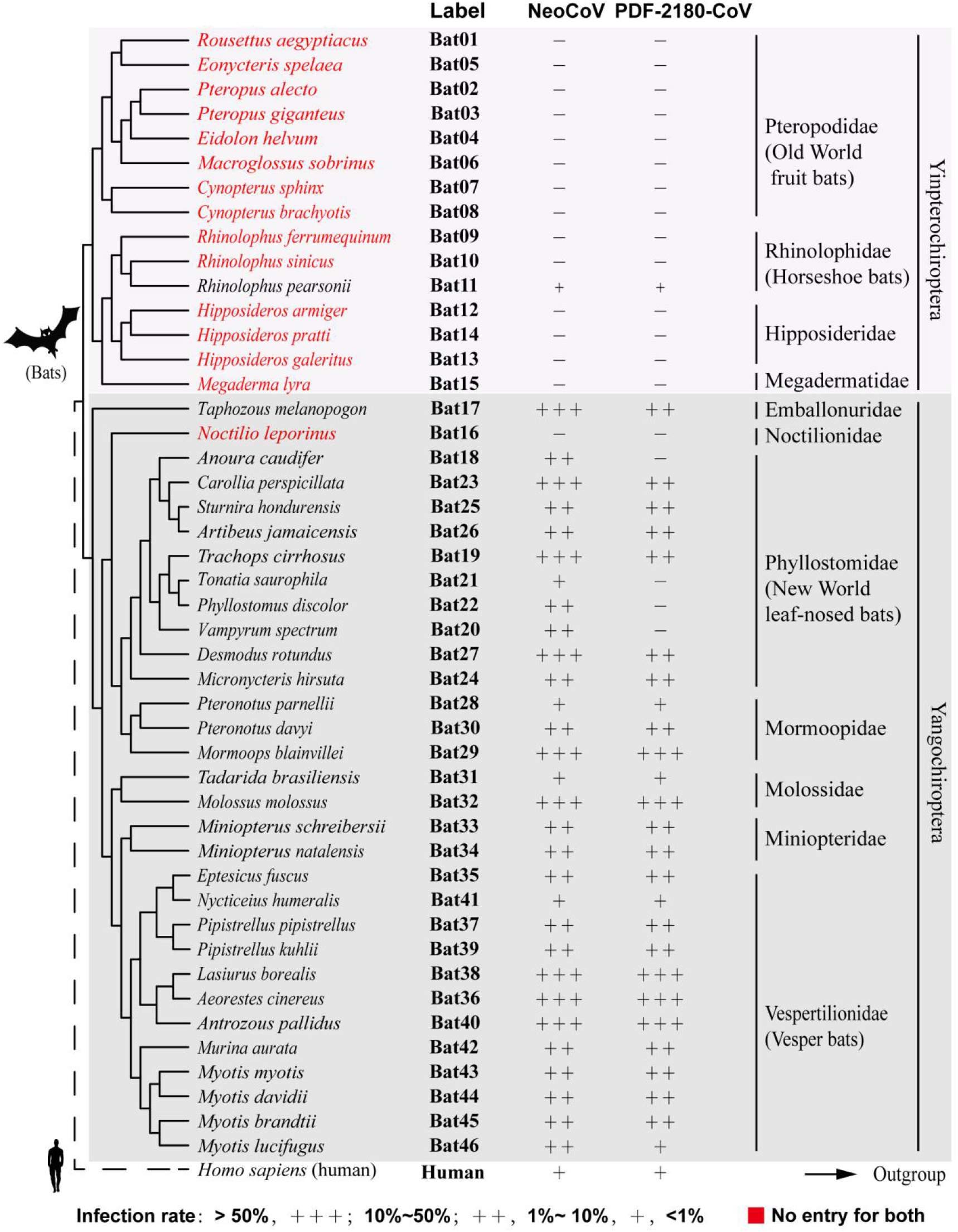
Receptor function of ACE2 from 46 bat species in supporting NeoCoV and PDF-2180-CoV entry.

**Extended Data Figure 3.**
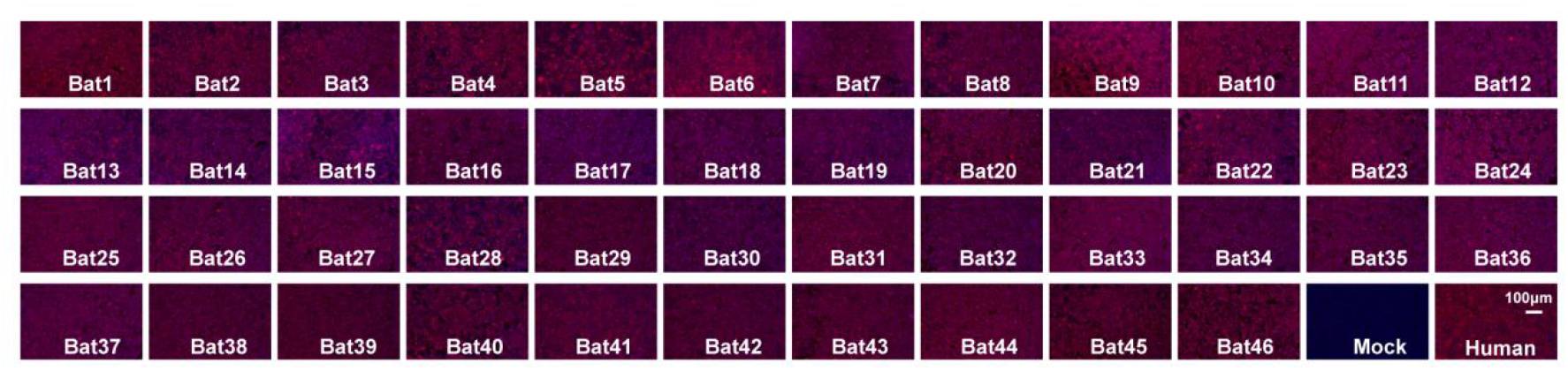
The expression level of 46 bat ACE2 orthologs in 293T cells as indicated by immunofluorescence assay detecting the C-terminal 3×FLAG Tag.

**Extended Data Figure 4.**
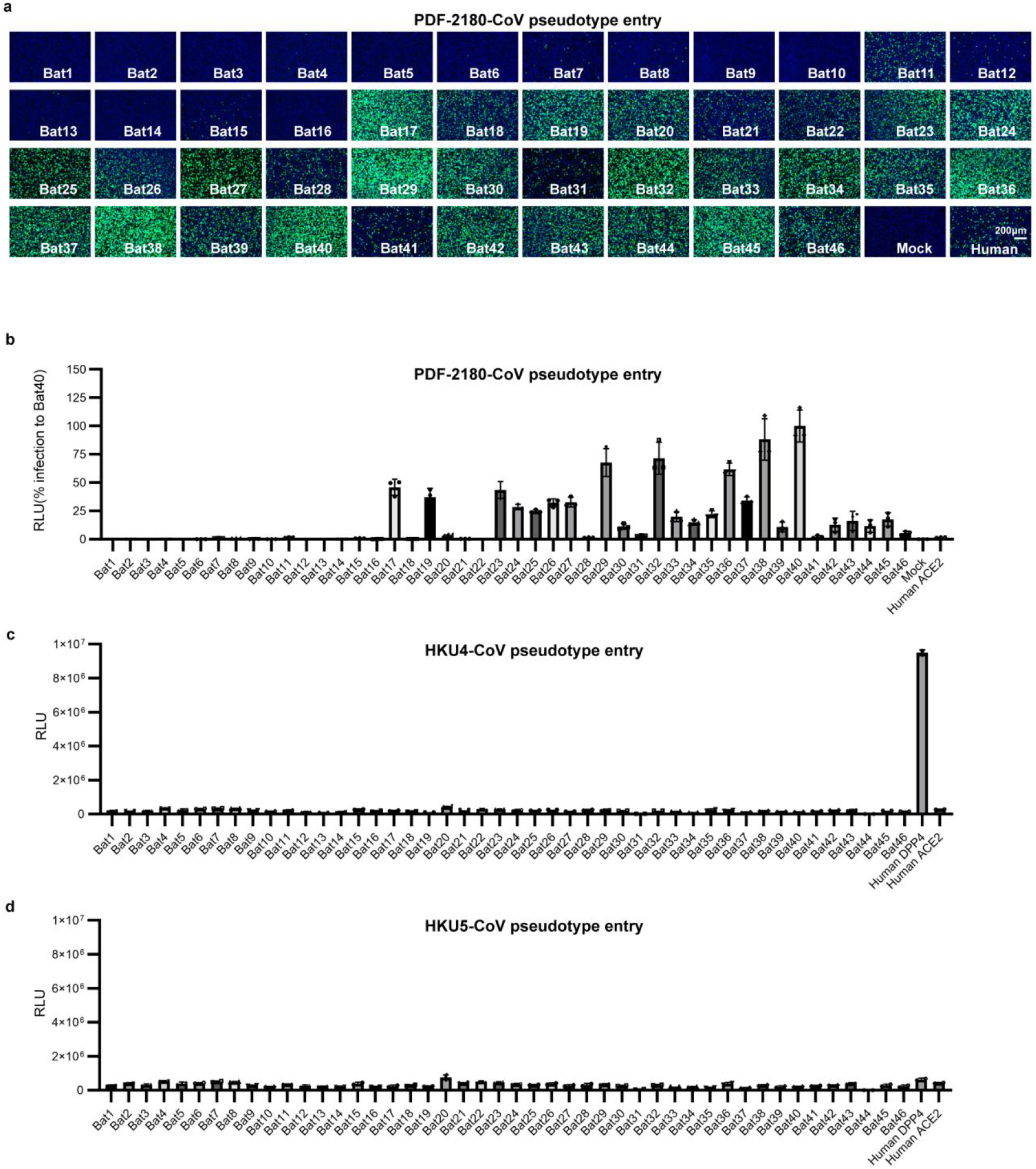
Entry efficiency of PDF-2180-CoV (a-b), HKU4-CoV (c), and HKU5-CoV (d) pseudoviruses in 293T cells expressing different bat ACE2 orthologs

**Extended Data Figure 5.**
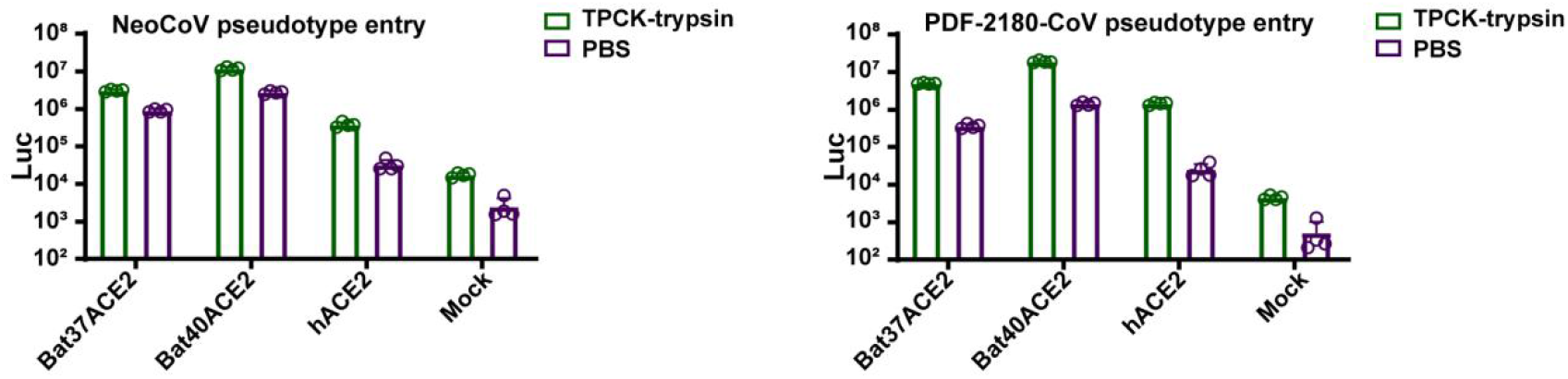
TPCK-trypsin treatment significantly boosted the entry efficiency of NeoCoV and PDF-2180-CoV on 293T cells expressing different ACE2 orthologs.

**Extended Data Figure 6.**
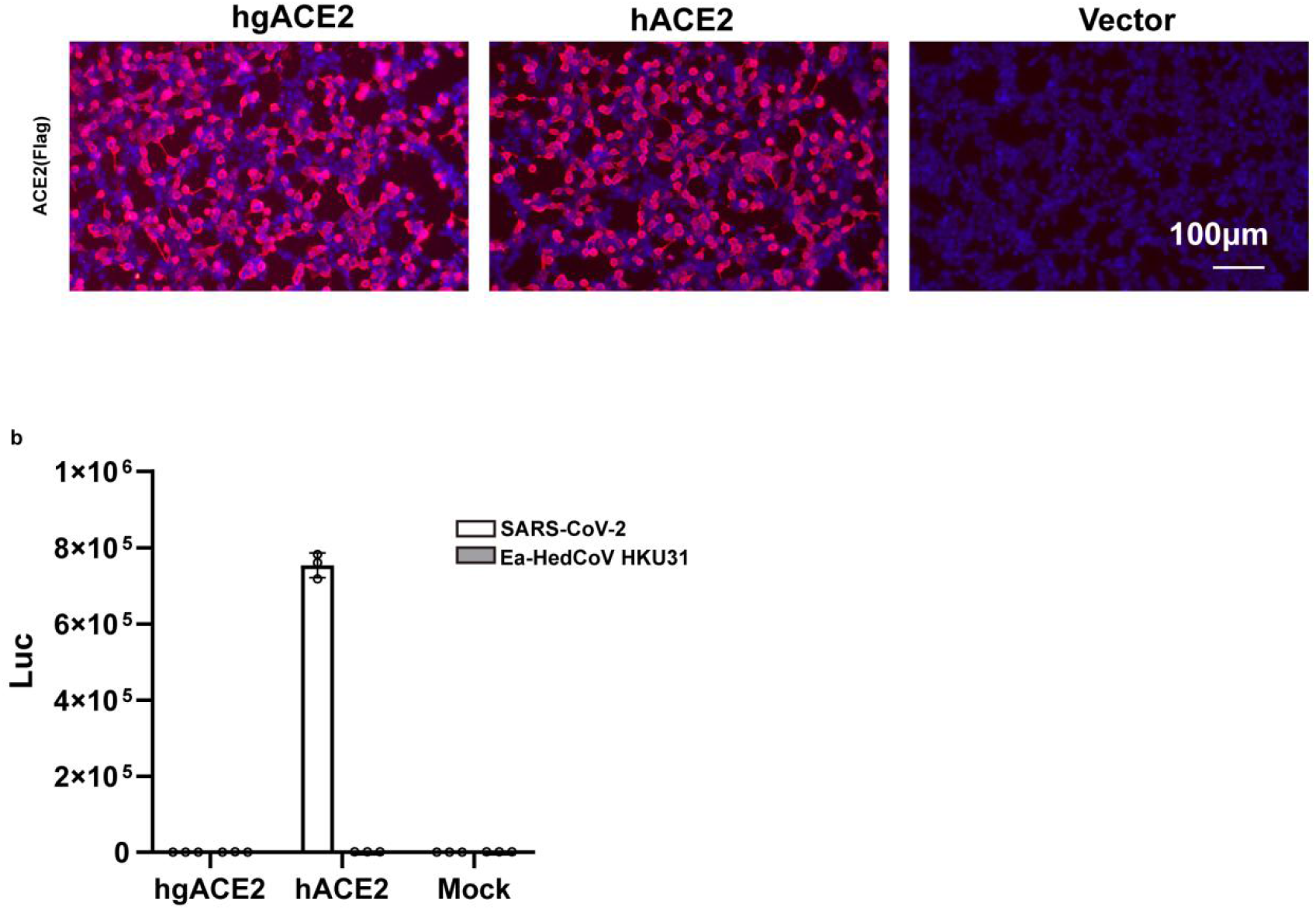
Hedgehog ACE2 (hgACE2) cannot support the entry of Ea-HedCoV-HKU31. (a) The expression level of ACE2 was evaluated by immunofluorescence detecting the C-terminal fused Flag tag. (b) Viral entry of SARS-CoV-2 and HKU31 into cells expressing hACE2 or hgACE2.

**Extended Data Figure 7.**
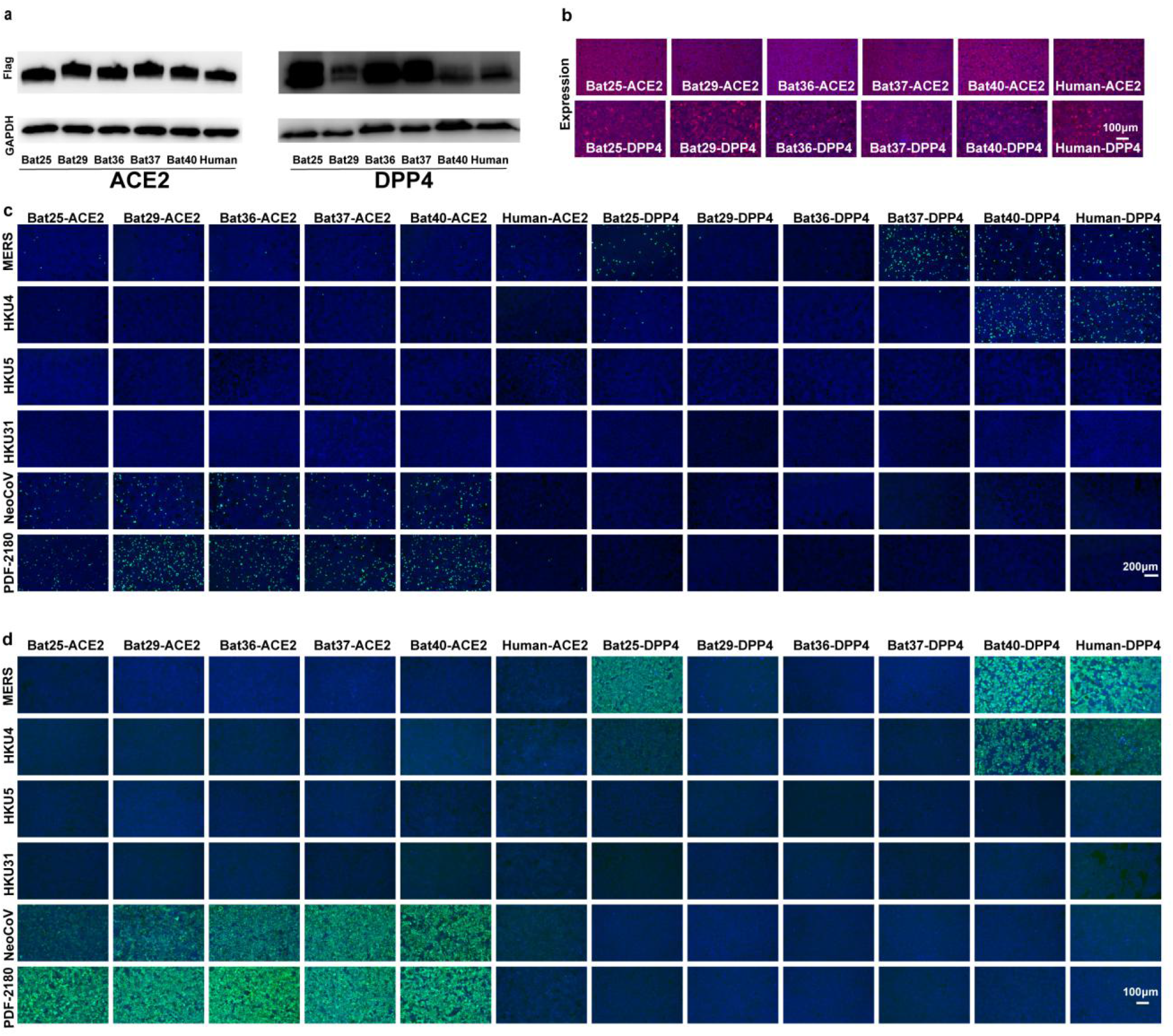
ACE2 and DPP4 receptor usage of different merbecoviruses. a, Western blot detected the expression levels of ACE2 and DPP4 orthologs in 293T cells.b, The intracellular bat ACE2 expression level by immunofluorescence assay detecting the C-terminal 3×FLAG-tag. c-d, Viral entry (c) and RBD binding (d) of different coronaviruses on 293T cells expressing different ACE2 and DPP4 orthologs.

**Extended data Figure 8.**
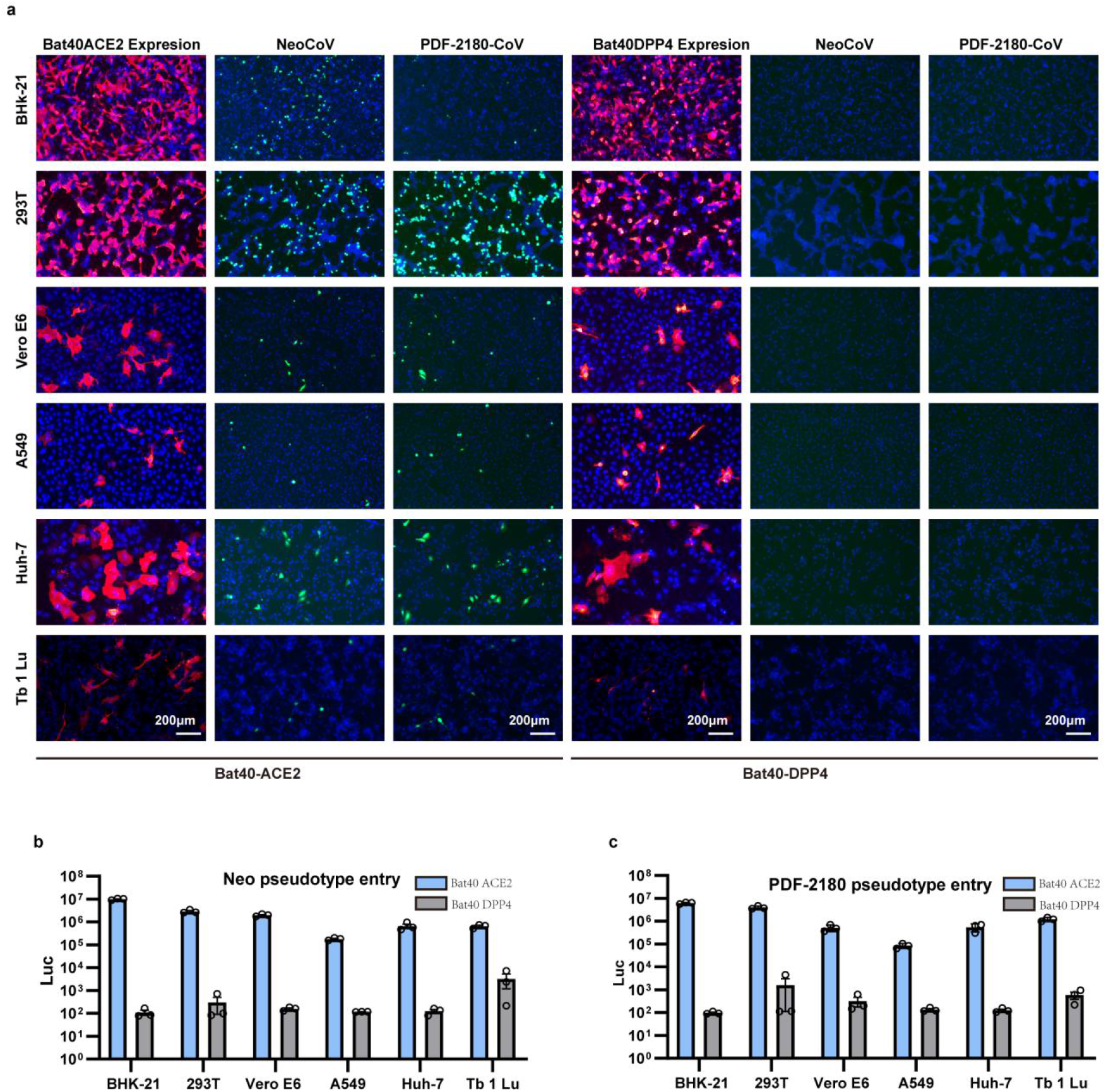
NeoCoV and PDF2180-CoV infection of different cell types expressing either Bat40ACE2 or Bat40DPP4. The BHK-21, 293T, Vero E6, A549, Huh-7, and Tb 1 Lu were transfected with either Bat40ACE2 or Bat40DPP4. The expression and viral entry (GFP) (a) and luciferase activity (c) were detected at 16 hpi.

**Extended data Figure 9.**
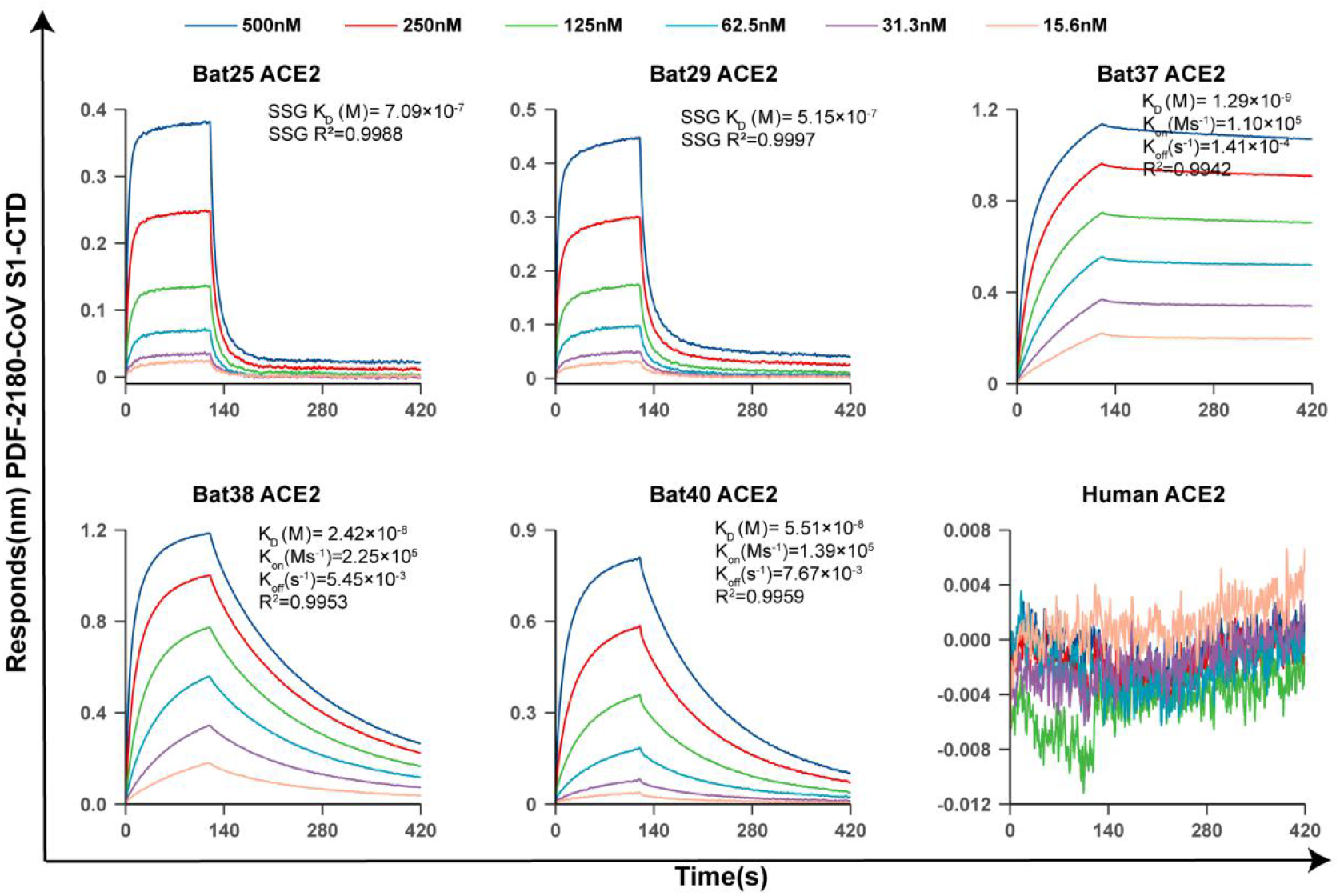
BLI analysis of the binding kinetics of PDF-2180-CoV S1-CTD interacting with different ACE2 orthologs.

**Extended data Figure 10.**
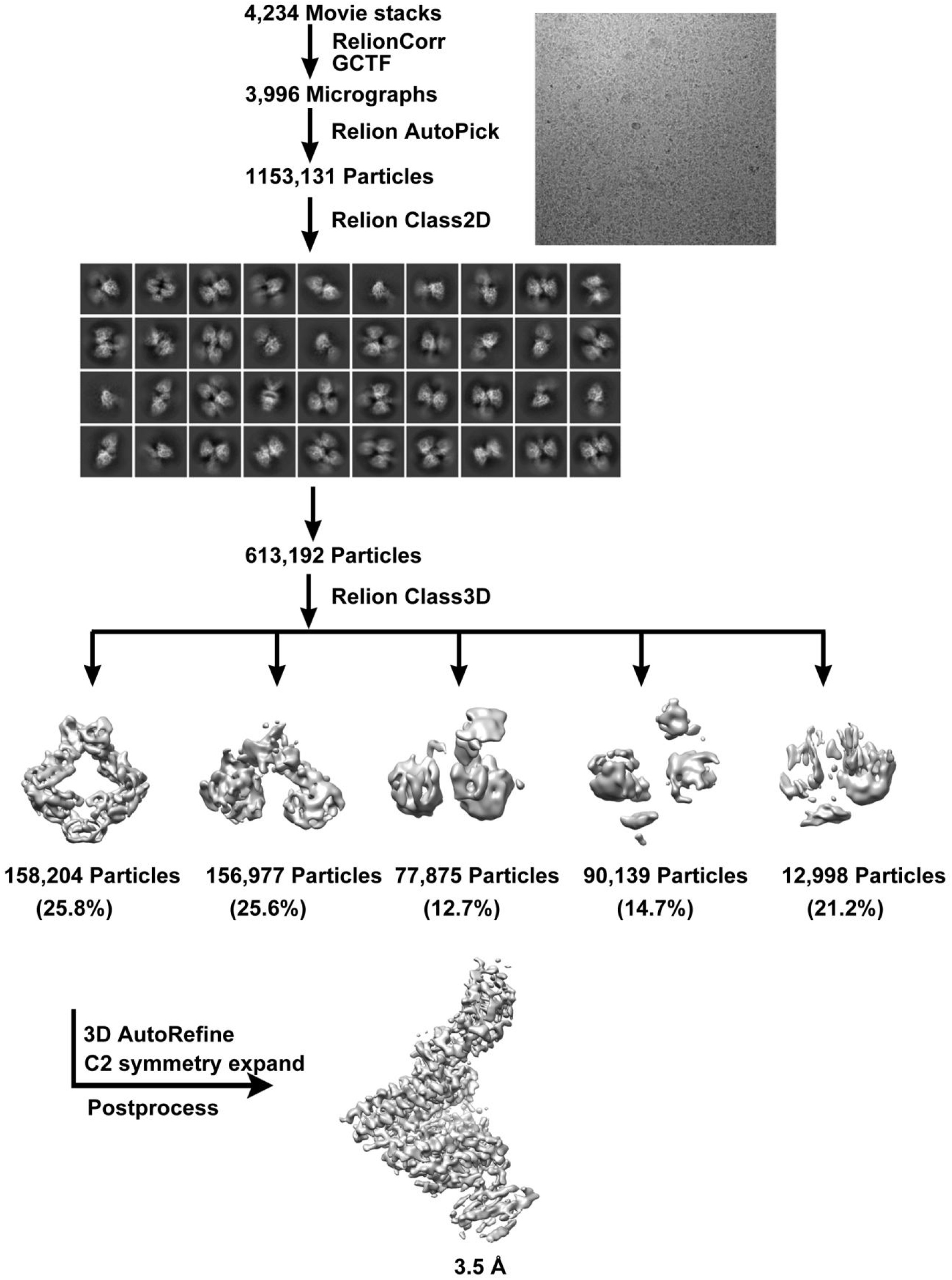
Flowcharts for cryo-EM data processing of Neo-CoV RBD-Bat37ACE2 complex.

**Extended data Figure 11.**
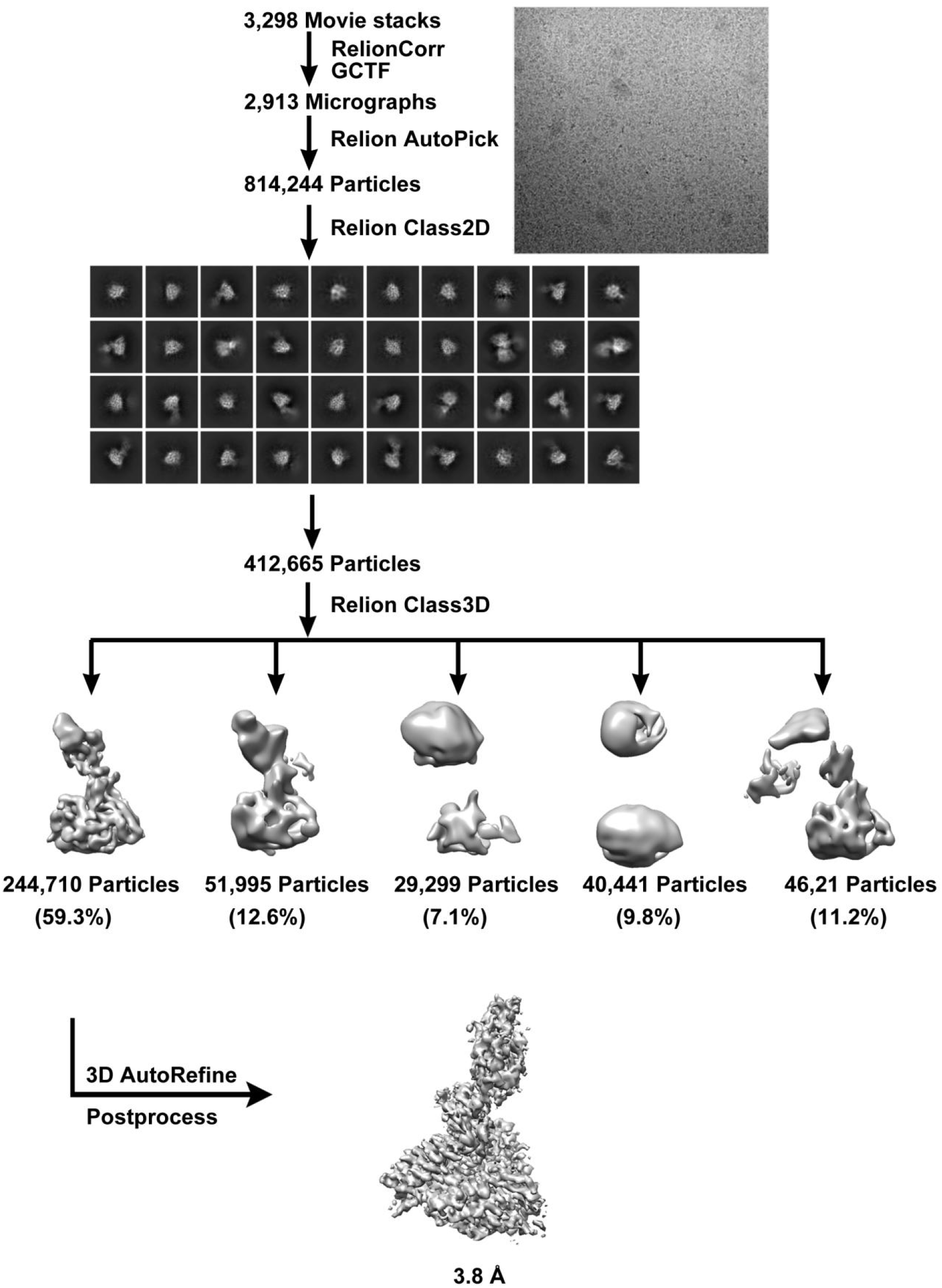
Flowcharts for cryo-EM data processing of PDF-2180-CoV RBD-Bat37ACE2 complex.

**Extended Data Figure 12.**
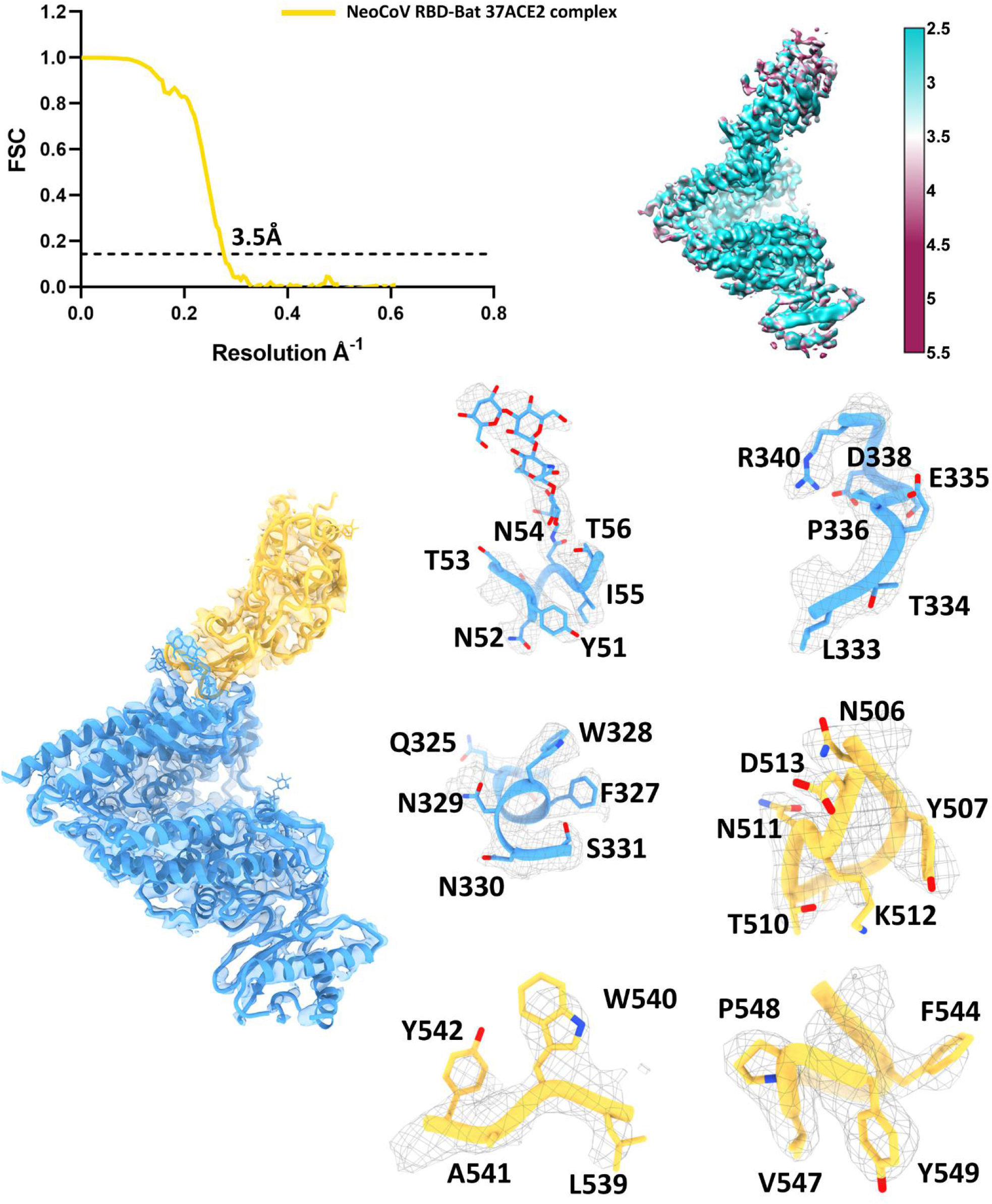
Resolution Estimation of the EM maps, density maps, and atomics models of NeoCoV RBD-Bat37ACE2 complex.

**Extended Data Figure 13.**
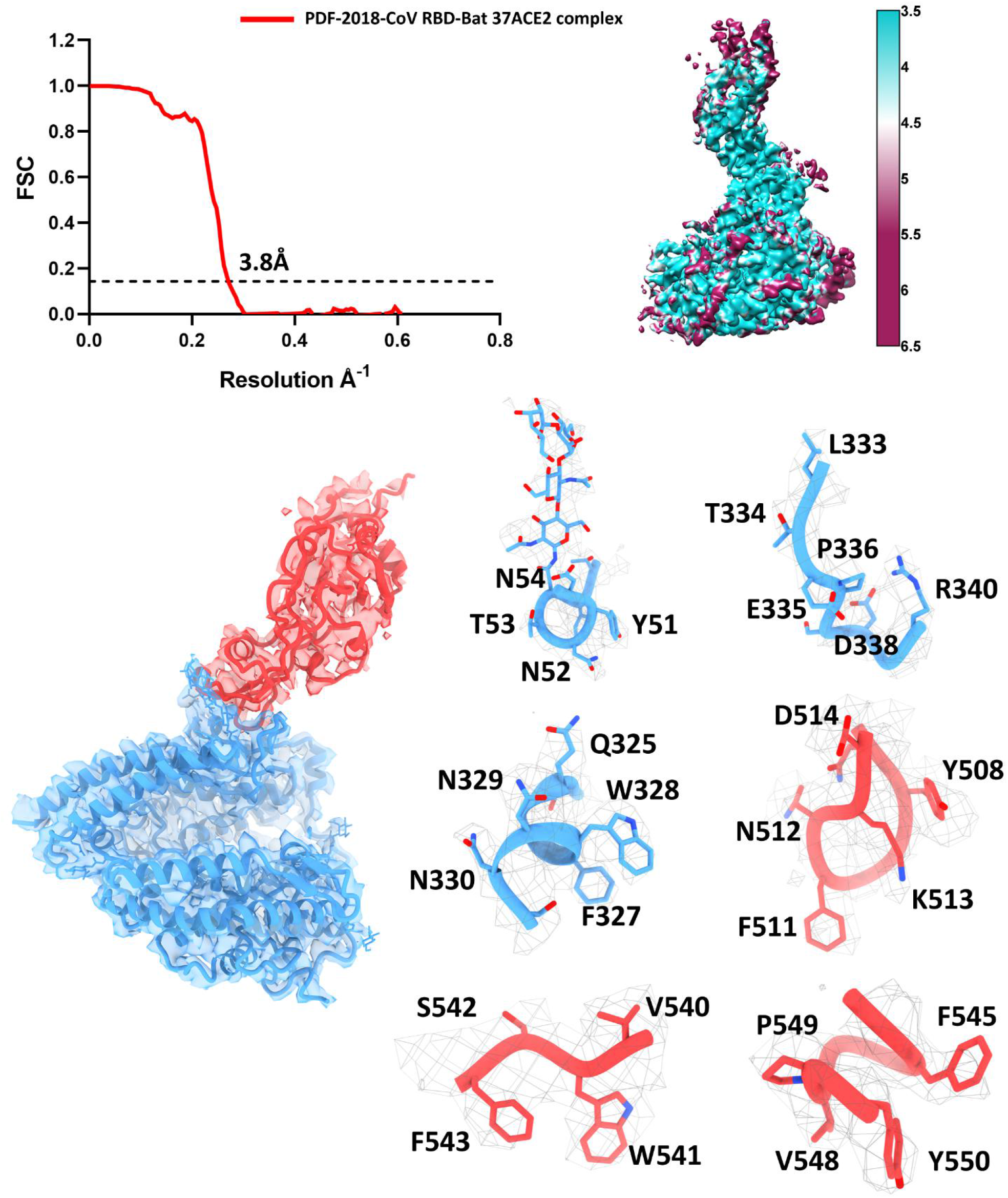
Resolution Estimation of the EM maps, density maps, and atomics models of PDF-2180-CoV RBD-Bat37ACE2 complex.

**Extended Data Figure 14.**
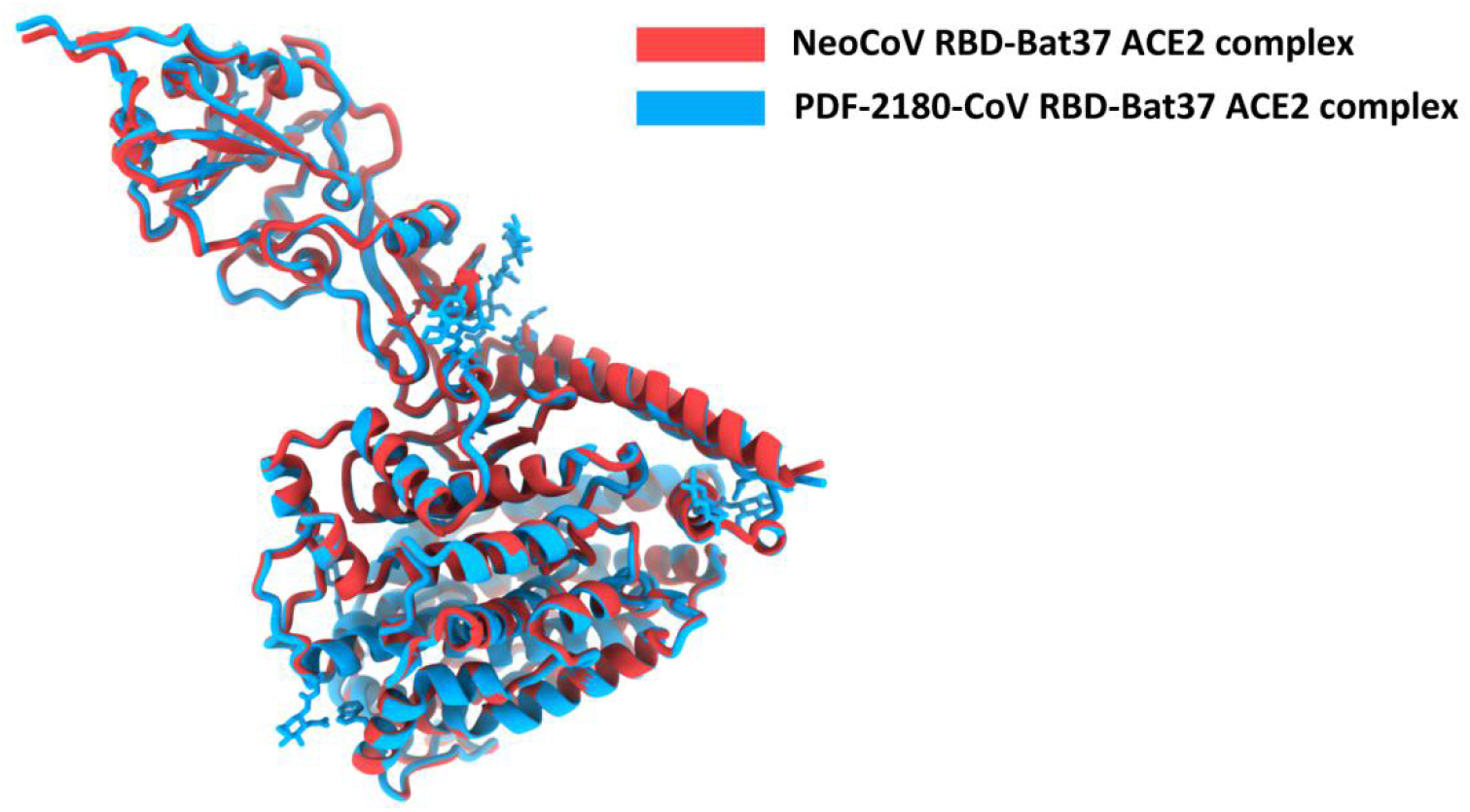
Superimposition of overall structures of NeoCoV RBD-Bat37ACE2 complex (red) and PDF-2018-COV RBD-Bat37ACE2 complex (bule).

**Extended Data Figure 15.**
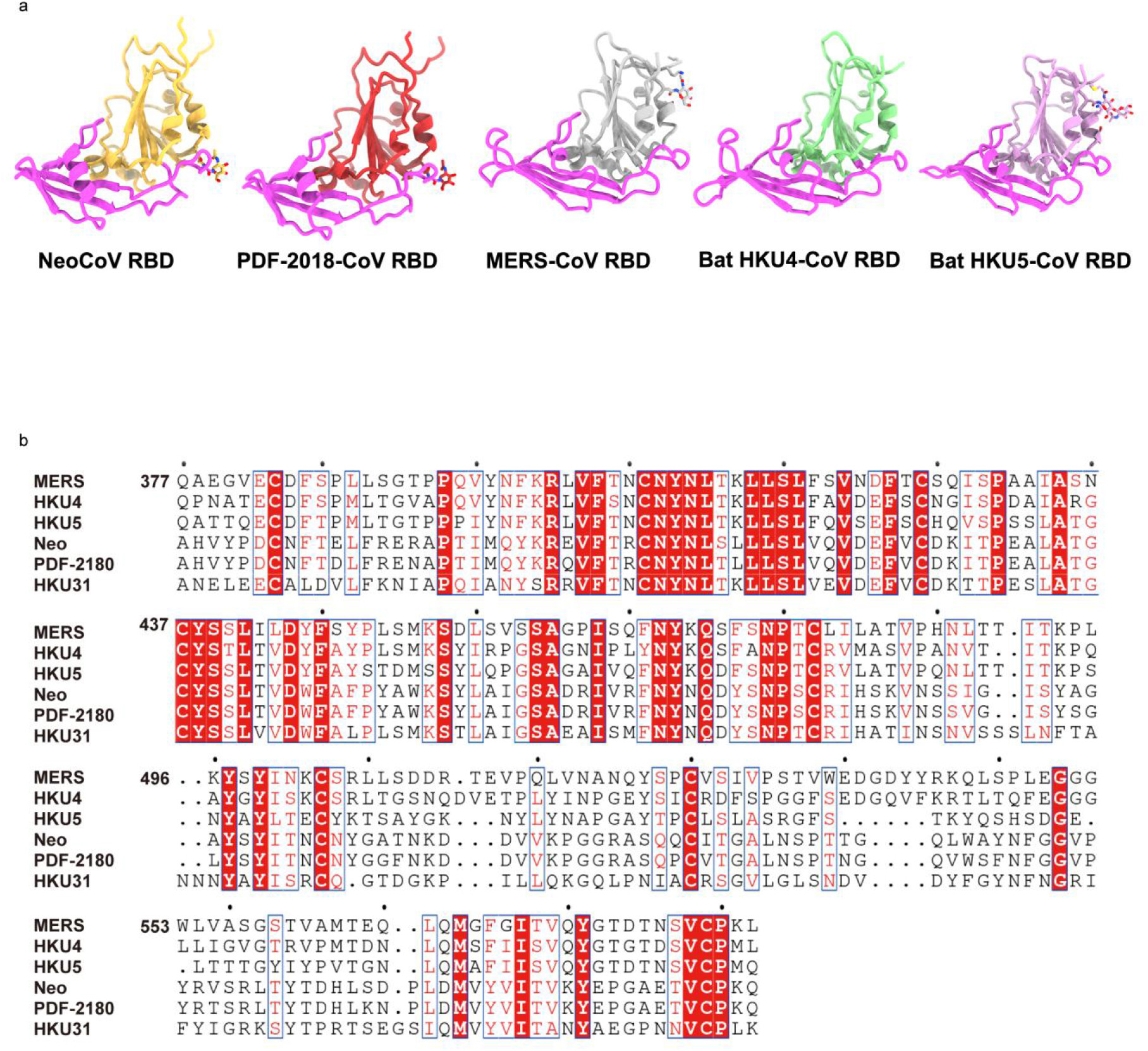
Structures and sequence comparison of RBDs from different merbecoviruses.

**Extended Data Figure 16.**
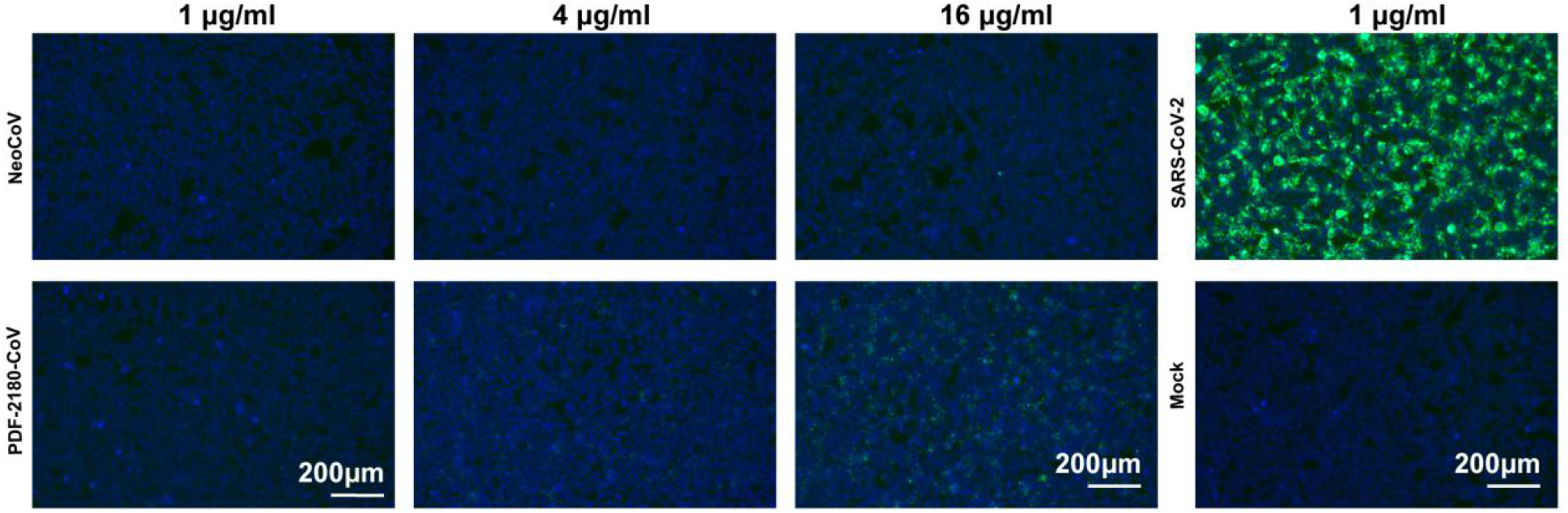
Comparison of the binding affinity of NeoCoV and PDF-2180-CoV RBD with hACE2 using SARS-CoV-2 RBD as a positive control.

**Extended Data Figure 17.**
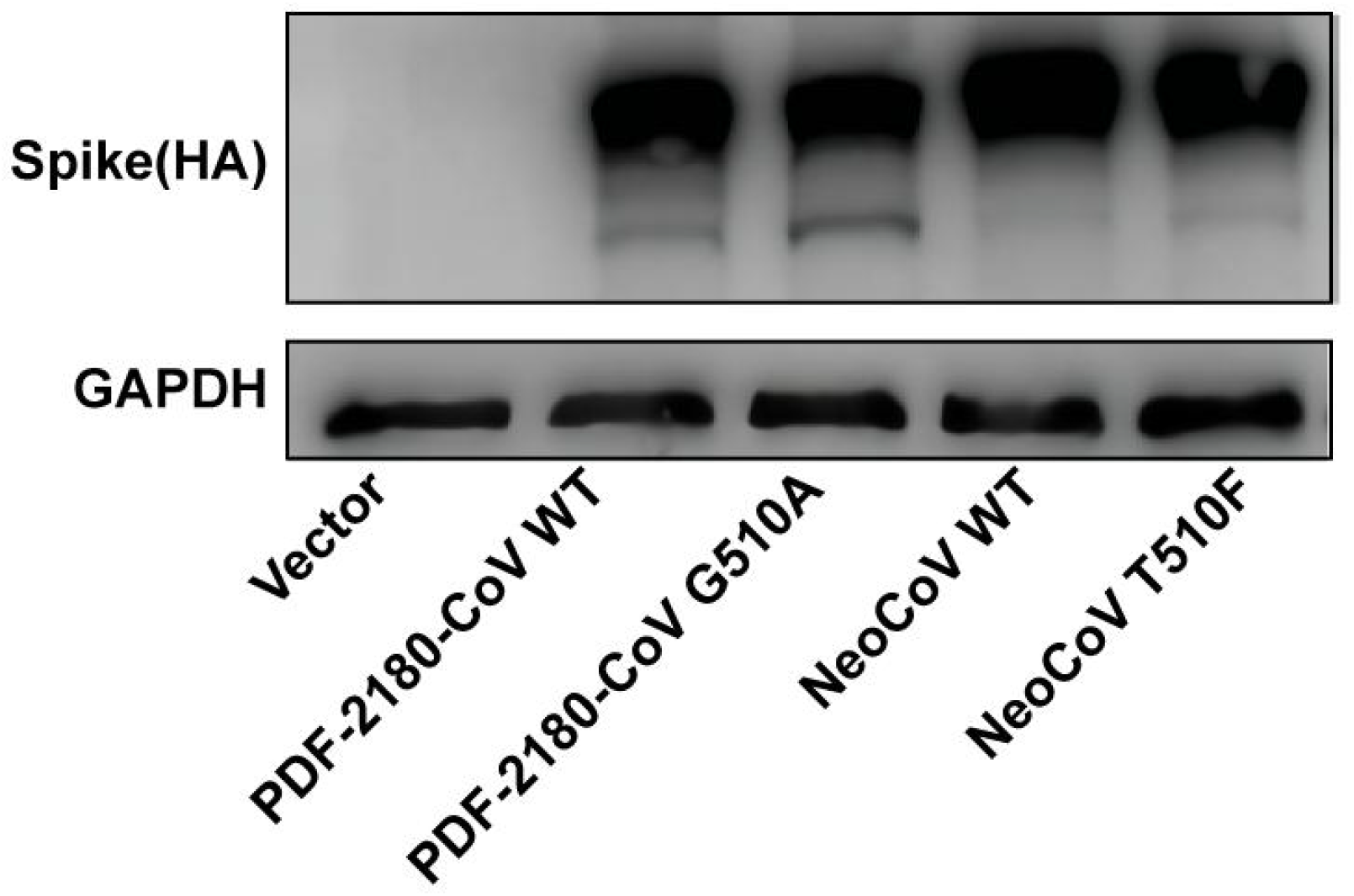
Expression level of the NeoCoV and PDF-2180-CoV spike proteins and their mutants.

**Extended Data Figure 18.**
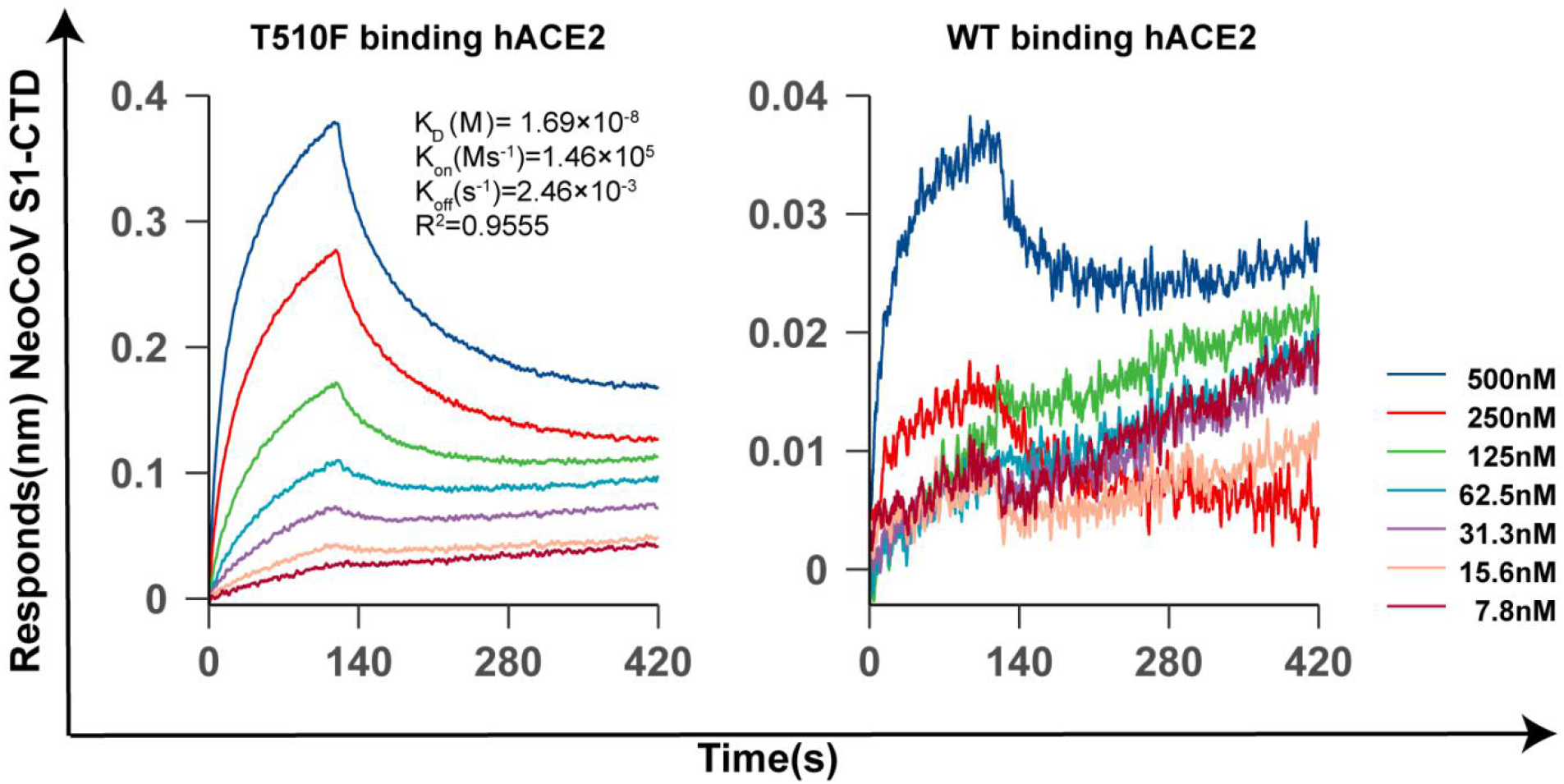
BLI analysis of the binding kinetics of NeoCoV S1-CTD WT and T510F interacting with human ACE2.

**Extended Data table 1.**
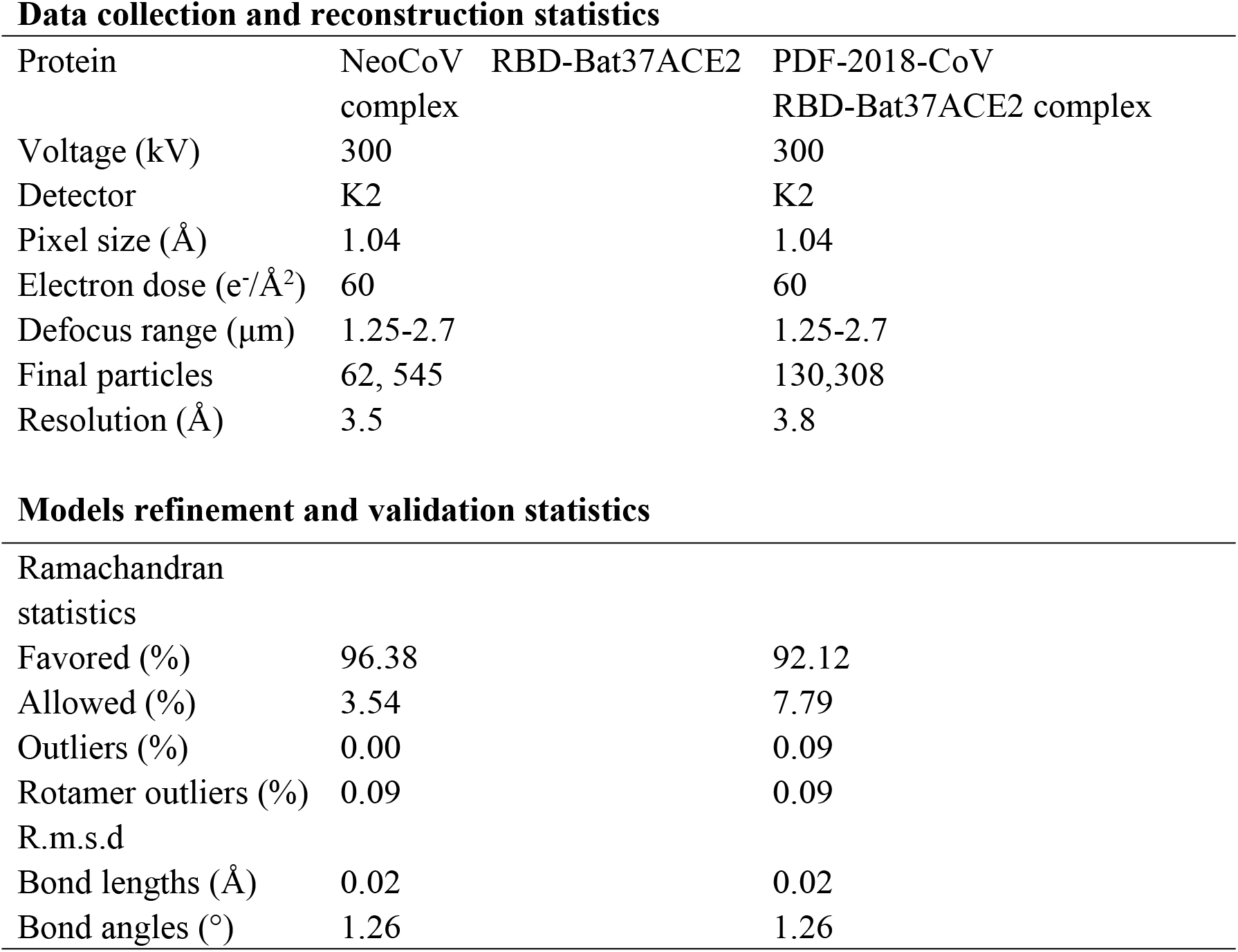
Cryo-EM data collection and atomic model refinement statistics of RBD mutant-mACE2 complex Data collection and reconstruction statistics

**Extended Data table 2.** Residues of NeoCoV RBD interacting with Bat37ACE2 at the binding interface (d < 4.5 Å)

| Bat37ACE2 |  | NeoCoV RBD |
| --- | --- | --- |
| Residues |  | Residues |
| E305 |  | T510 |
|  |  | K512 |
| W328 |  | A509 |
| N329 |  | G508 |
|  |  | A509 |
|  |  | G546 |
|  |  | P548 |
| S331 |  | A509 |
| M332 |  | A509 |
| L333 |  | A509 |
|  |  | T510 |
| T334 |  | A509 |
|  |  | T510 |
|  |  | N511 |
| E335 |  | N511 |
| P336 |  | N511 |
| D338 |  | N504 |
|  |  | N506 |
|  |  | L539 |
| R340 |  | N511 |
|  |  | R550 |

**Extended Data table 3.** Residues of PDF-2180-CoV RBD interacting with Bat37ACE2 at the binding interface (d < 4.5 Å)

| **Bat37ACE2** | **PDF-2180-CoV RBD** |
| --- | --- |
| **Residues** | **Residues** |
| A304 | F511 |
| E305 | F511 |
|  | K513 |
| F308 | F511 |
| W328 | G510 |
|  | F511 |
| N329 | G509 |
|  | G510 |
|  | G547 |
|  | P549 |
| N330 | N544 |
|  | G547 |
|  | P549 |
| S331 | G510 |
| M332 | G510 |
| L333 | G510 |
|  | F511 |
| T334 | F511 |
|  | N512 |
| E335 | N512 |
| P336 | N512 |
| D338 | N505 |
|  | N507 |
|  | L540 |
| R340 | N512 |
|  | R551 |

**Extended Data table 4.** MD prediction of the effect of critical residue variations to the interaction between NeoCoV and Bat37ACE2 by mCSM-PPI2.

| Substitution | $\Delta \Delta G$ (kcal/mol) | | | | | | | | | |
| --- | --- | --- | --- | --- | --- | --- | --- | --- | --- | --- |
|  | N504 | G508 | A509 | T510 | N511 | K512 | L539 | G546 | P548 | R550 |
| A | -0.087 | -0.573 |  | 0.041 | -0.477 | -0.396 | -0.22 | -0.106 | -0.646 | -0.434 |
| R | 0.133 | -0.849 | 0.261 | 0.196 | -0.315 | 0.001 | -0.273 | -0.164 | -0.152 |  |
| N |  | -0.515 | 0.466 | 0.078 |  | 0.015 | -0.283 | -0.008 | -0.33 | -0.369 |
| D | 0.098 | -0.401 | 0.518 | 0.179 | -0.038 | -0.362 | -0.257 | 0.219 | -0.167 | -0.543 |
| C | -0.103 | -0.805 | 0.148 | -0.052 | -0.525 | -0.473 | -0.261 | -0.223 | -0.385 | -0.405 |
| Q | 0.113 | -0.572 | 0.341 | 0.077 | -0.577 | 0.211 | -0.215 | -0.007 | -0.121 | -0.285 |
| E | 0.294 | -0.324 | 0.37 | 0.131 | -0.277 | -0.348 | -0.302 | 0.21 | 0.006 | -0.486 |
| G | -0.129 |  | -0.671 | -0.102 | -0.435 | -0.543 | -0.236 |  | 0.147 | -0.591 |
| H | 0.156 | -0.64 | 0.402 | 0.146 | -0.065 | -0.157 | -0.064 | -0.16 | -0.304 | -0.12 |
| I | -0.182 | -0.787 | 0.403 | 0.081 | -0.184 | -0.319 | -0.178 | -0.177 | -0.403 | -0.265 |
| L | -0.137 | -0.857 | 0.334 | 0.061 | -0.123 | -0.02 |  | -0.189 | -0.508 | -0.269 |
| K | 0.27 | -0.767 | 0.009 | 0.416 | -0.722 |  | -0.175 | -0.098 | -0.187 | -0.113 |
| M | -0.255 | -0.682 | -0.1 | -0.001 | -0.246 | -0.325 | -0.251 | -0.105 | -0.526 | -0.311 |
| F | -0.073 | -0.476 | 1.012 | 0.487 | -0.12 | -0.134 | 0.287 | -0.023 | 0.1504 | -0.281 |
| P | 0.051 | -0.712 | -0.44 | -0.149 | -0.478 | -0.41 | -0.379 | -0.125 |  | -0.161 |
| S | -0.101 | -0.441 | 0.237 | -0.011 | -0.454 | -0.17 | -0.142 | -0.082 | -0.279 | -0.3 |
| T | -0.061 | -0.567 | 0.342 |  | -0.476 | -0.226 | -0.186 | -0.077 | -0.414 | -0.32 |
| W | -0.002 | -0.158 | 1.396 | 0.481 | 0.033 | -0.132 | 0.234 | 0.09 | 0.335 | -0.066 |
| Y | -0.072 | -0.439 | 1.355 | 0.284 | -0.083 | -0.065 | 0.204 | 0.02 | 0.1918 | -0.282 |
| V | -0.086 | -0.673 | 0.025 | 0.003 | -0.341 | -0.408 | 0.031 | -0.082 | -0.563 | -0.405 |

